# Ascending neurons convey behavioral state to integrative sensory and action selection centers in the brain

**DOI:** 10.1101/2022.02.09.479566

**Authors:** Chin-Lin Chen, Florian Aymanns, Ryo Minegishi, Victor D. V. Matsuda, Nicolas Talabot, Semih Günel, Barry J. Dickson, Pavan Ramdya

## Abstract

Knowledge of one’s own behavioral state—whether one is walking, grooming, or resting—is critical for contextualizing sensory cues including interpreting visual motion and tracking odor sources. Additionally, awareness of one’s own posture is important to avoid initiating destabilizing or physically impossible actions. Ascending neurons (ANs), interneurons in the vertebrate spinal cord or insect ventral nerve cord (VNC) that project to the brain, may provide such high-fidelity behavioral state signals. However, little is known about what ANs encode and where they convey signals in any brain. To address this gap, we performed a large-scale functional screen of AN movement encoding, brain targeting, and motor system patterning in the adult fly, *Drosophila melanogaster*. Using a new library of AN sparse driver lines, we measured the functional properties of 247 genetically-identifiable ANs by performing two-photon microscopy recordings of neural activity in tethered, behaving flies. Quantitative, deep network-based neural and behavioral analyses revealed that ANs nearly exclusively encode high-level behaviors—primarily walking as well as resting and grooming—rather than low-level joint or limb movements. ANs that convey self-motion—resting, walking, and responses to gust-like puff stimuli—project to the brain’s anterior ventrolateral protocerebrum (AVLP), a multimodal, integrative sensory hub, while those that encode discrete actions—eye grooming, turning, and proboscis extension—project to the brain’s gnathal ganglion (GNG), a locus for action selection. The structure and polarity of AN projections within the VNC are predictive of their functional encoding and imply that ANs participate in motor computations while also relaying state signals to the brain. Illustrative of this are ANs that temporally integrate proboscis extensions over tens-of-seconds, likely through recurrent interconnectivity. Thus, in line with long-held theoretical predictions, ascending populations convey high-level behavioral state signals almost exclusively to brain regions implicated in sensory feature contextualization and action selection.

## 1 Introduction

To generate adaptive behaviors, animals [1] and robots [2] must not only sense their environment but also be aware of their own behavioral state including low-level movements of their limbs and high-level behaviors such as walking and resting. This self-awareness has long been theorized to overcome at least two major challenges for robust, autonomous control. First, knowing if one is at rest or in motion permits the accurate interpretation of whether sensory cues, like visual motion during feature tracking or odor intensity fluctuations during plume following, result from exafference (the movement of objects in the world), or reafference (self-motion with respect to stationary objects) [1]. Second, being aware of one’s current posture enables the selection of appropriate future actions that are not destabilizing, or physically impossible.

In line with these theoretical predictions, neural representations of behaviors have been observed widely across the brains of mice [3–5], and in the fly, *Drosophila melanogaster* [6–9]. Furthermore, studies in *Drosophila* have supported roles for behavioral state signals in sensory contextualization (flight [6] and walking [7] modulate neurons in the visual system [8, 10]), and action selection (an animal’s walking speed regulates its decision to run or freeze in response to a fear-inducing stimulus [11]).

Despite these advances, the cellular origins of behavioral state signals in the brain remain largely unknown. On one hand, they might arise from efference copies generated by descending neurons (DNs) in the brain that project to and drive downstream motor systems [1]. However, these efference copies would not be expected to provide the most precise readout of one’s own behavioral state: the brain’s descending commands will be sculpted by musculoskeletal interactions with the environment. Instead, a more categorically and temporally precise readout of ongoing behaviors might be obtained from ascending neurons in the motor system that process proprioceptive and tactile signals and then convey a holistic representation of behavioral states to the brain. Although these behavioral signals may come from a subset of primary mechanosensory neurons in the limbs [12], they are more likely to be computed and conveyed by second- and higher-order ascending neurons (ANs) residing in the spinal cord of vertebrates [13–16], or insect ventral nerve cord (VNC) [17, 18]. In *Drosophila*, ANs have been shown to process limb proprioceptive and tactile signals, likely sculpting a more complex and ethologically-salient readout of ongoing movements [12, 19, 20].

To date only a few genetically-identifiable AN cell types have been studied in behaving animals— primarily in the fly, *Drosophila melanogaster*, which has a relatively small number of neurons that can also be genetically targeted for repeated investigation. These studies support the hypothesis that ANs are a prominent source of behavioral state signals in the brain. First, microscopy recordings of AN terminals in the brain have shown that Lco2N1 and Les2N1D ANs are active during walking [21], and that LAL-PS-ANs convey walking signals to the visual system [22]. Second, artificial activation of pairs of PER_in_ ANs [23], and Moonwalker ANs [24] regulates action selection and behavioral persistence, respectively.

These first insights urgently motivate the investigation of three fundamental questions via a more comprehensive and quantitative analysis of large AN populations. First, what information do ANs convey to the brain **(Figure S1A)**? They might encode low-level movements of the joints or limbs, or high-level behavioral states like whether an animal is walking, or grooming **(Figure 1A-i)**. Second, where do ANs convey this information to in the brain **(Figure S1B)**? They might project widely across brain regions, or narrowly target circuit hubs with specific functions **(Figure 1A-ii)**. Third, what can an AN’s patterning within the VNC tell us about its encoding and computational role **(Figure 1A-iii)**? Answering these three questions would open the door to a cellular-level understanding of how neurons encode behavioral states by integrating proprioceptive, tactile, and other sensory feedback signals. It would also enable the study of how behavioral state signals are incorporated by brain circuits to intelligently contextualize multimodal cues and to select appropriate future actions.

**Figure 1:**
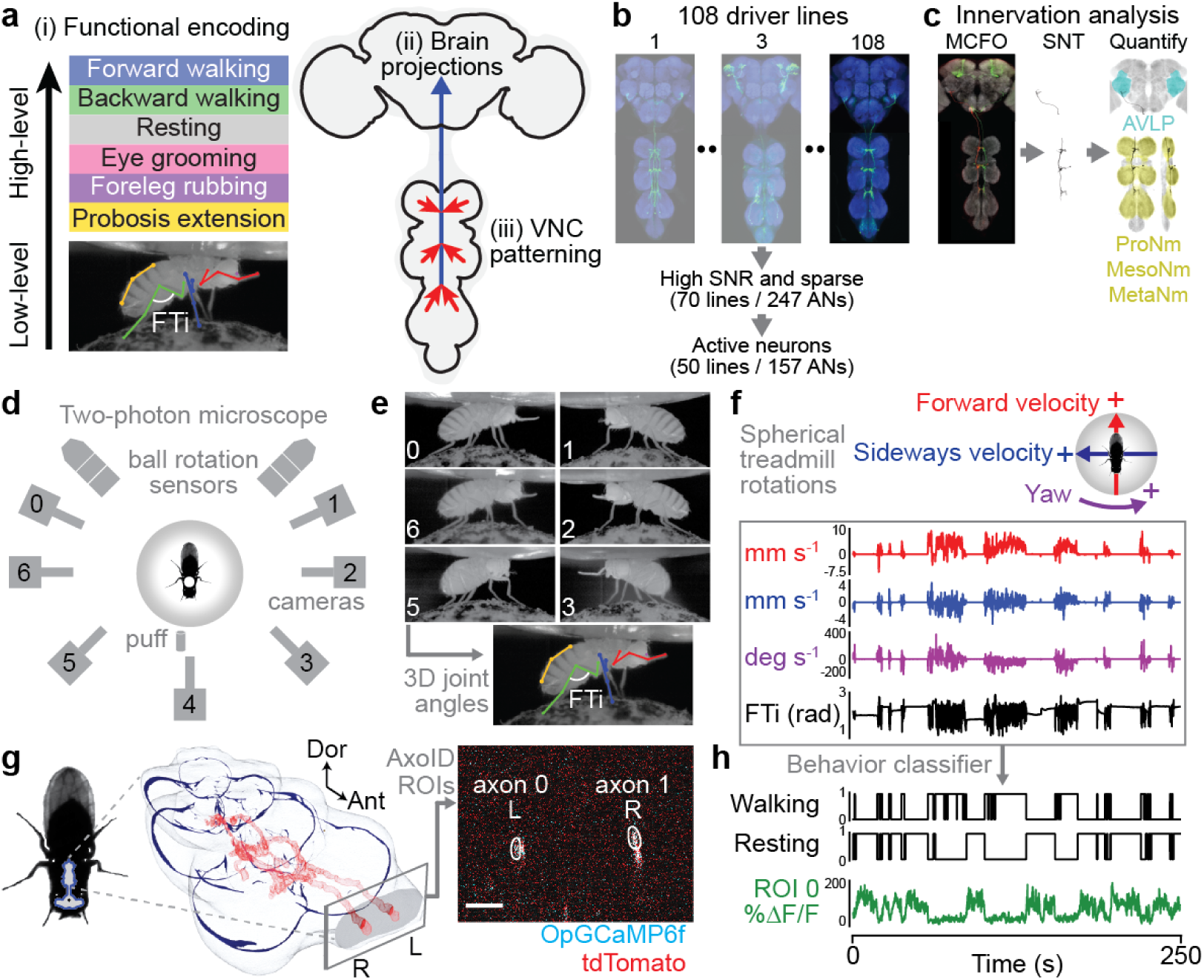
Large-scale functional and morphological screen of ascending neuron movement encoding and nervous system targeting. **(a)** Schematic of the main questions addressed. (i) To what extent do ascending neurons (ANs) encode high-level behaviors, or low-level movements? (ii) Where in the brain do ANs convey behavioral states? (iii) To what extent is an AN’s patterning within the VNC predictive of its encoding? **(b)** Screening 108 driver lines. The projection patterns of sparse lines with active ANs and high SNR (157 ANs) were examined in the brain and VNC. **(c)** These were quantified using broad spFP and single-cell MCFO confocal imaging. **(d)** Overhead schematic of the behavior measurement system used during two-photon microscopy. A camera array captures six views of the animal. Two optic flow sensors measure ball rotations. A puff of CO_2_ (or air) is used to elicit behavior from sedentary animals. **(e)** 2D poses are estimated for six cameras views using DeepFly3D. These data are triangulated to quantify 3D poses and joint angles for six legs and the abdomen (color-coded). The Femur-Tibia (FTi) joint angle is indicated (white). **(f)** Two optic flow sensors measure rotations of the spherical treadmill as a proxy for forward (red), sideways (blue), and yaw (purple) walking velocities. Positive directions of rotation (‘+’) are indicated. **(g, left)** A volumetric representation of the ventral nerve cord (VNC) including a reconstruction of ANs targeted by the SS27485-spGal4 driver (red). Indicated are the dorsal-ventral (‘Dor’) and anterior-posterior (‘Ant’) axes, as well as the fly’s left (L) and right (R) sides. **(g, right)** Sample two-photon cross-section image of the thoracic neck connective showing ANs that express OpGCaMP6f (cyan) and tdTomato (red). AxoID is used to semi-automatically identify two axonal regions-of-interest (ROIs, white) on the left (‘L’) and right (‘R’) sides of the connective. **(h)** Spherical treadmill rotations and joint angles are used to classify behaviors. Binary classifications are then compared with simultaneously recorded neural activity for 250 s trials of spontaneous and puff-elicited behaviors. Shown is an activity trace from ROI 0 (green) in panel **g**.

To address these questions, we developed and used a number of advanced experimental and analytical tools. First, we screened a library of split-Gal4 *Drosophila* driver lines (R.M. and B.J.D., unpublished). These, along with the published MAN-spGal4 [24] and 12 sparsely expressing Gal4 lines [25], collectively allowed us to gain repeated genetic access to 247 ANs (**Figure 1B**; **Table 1**). Using these driver lines and a multi-color flip-out (MCFO) approach [26], we then quantified the projections of ANs within the brain and VNC (**Figure 1C**). Second, we screened the encoding of these ANs through two-photon microscopy functional recordings of neural activity within the VNC of tethered, behaving flies [27]. To overcome noise and movement-related deformations in imaging data, we developed and used ‘AxoID’, a deep learning-based software to semi-automatically identify and track axonal Regions-of-Interest (ROIs)(see Methods). Third, to precisely quantify joint angles and limb kinematics, we used a multicamera array to record behavior during two-photon imaging. We processed resulting videos using DeepFly3D, a deep learning-based 3D pose estimation software [28]. By combining these 3D joint positions with measured spherical treadmill rotations, a proxy for locomotor velocities [29], we could then segment and classify behavioral time-series and study the relationship between behavioral states and ongoing neural activity using linear models.

**Table 1:**
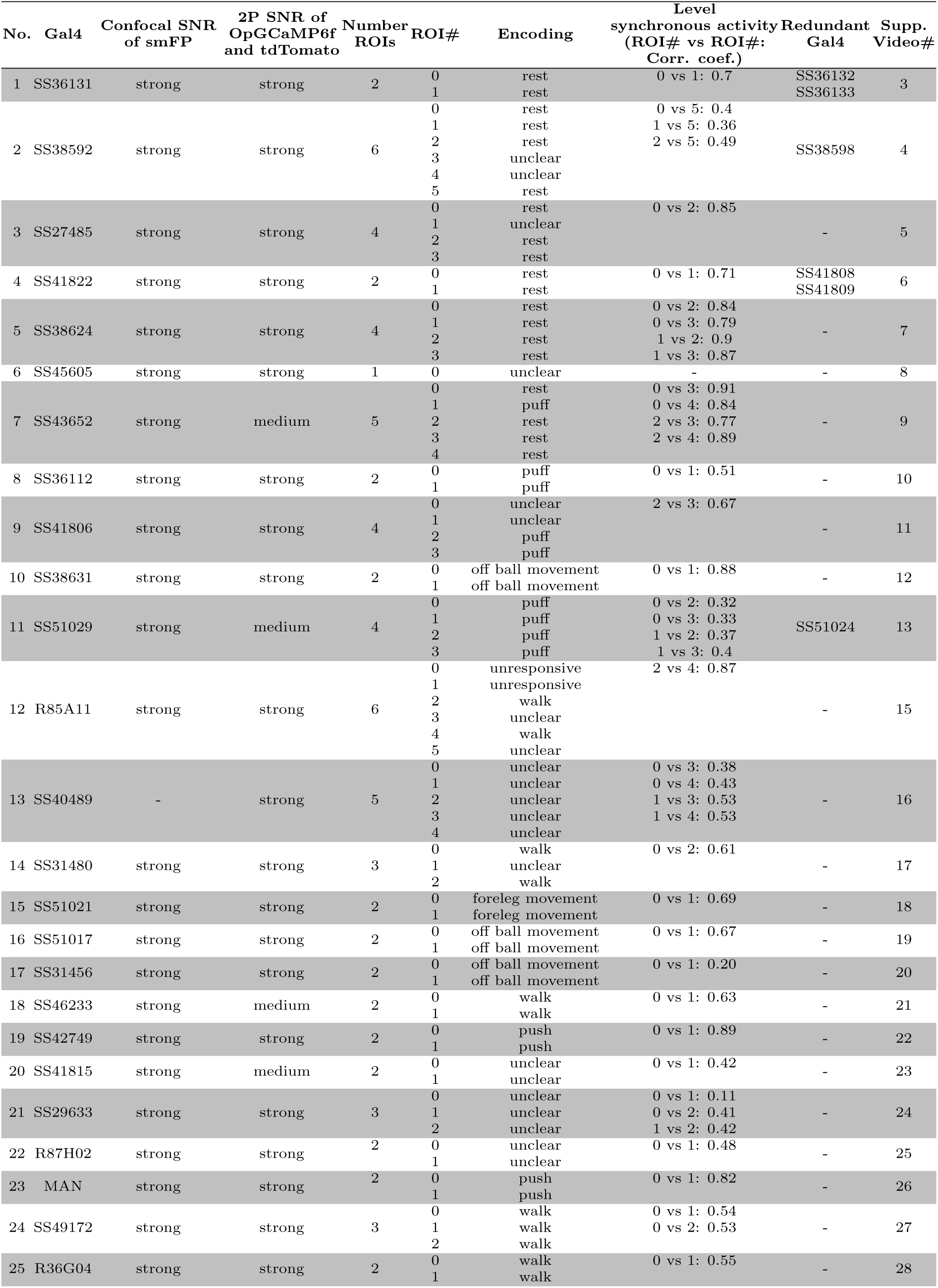
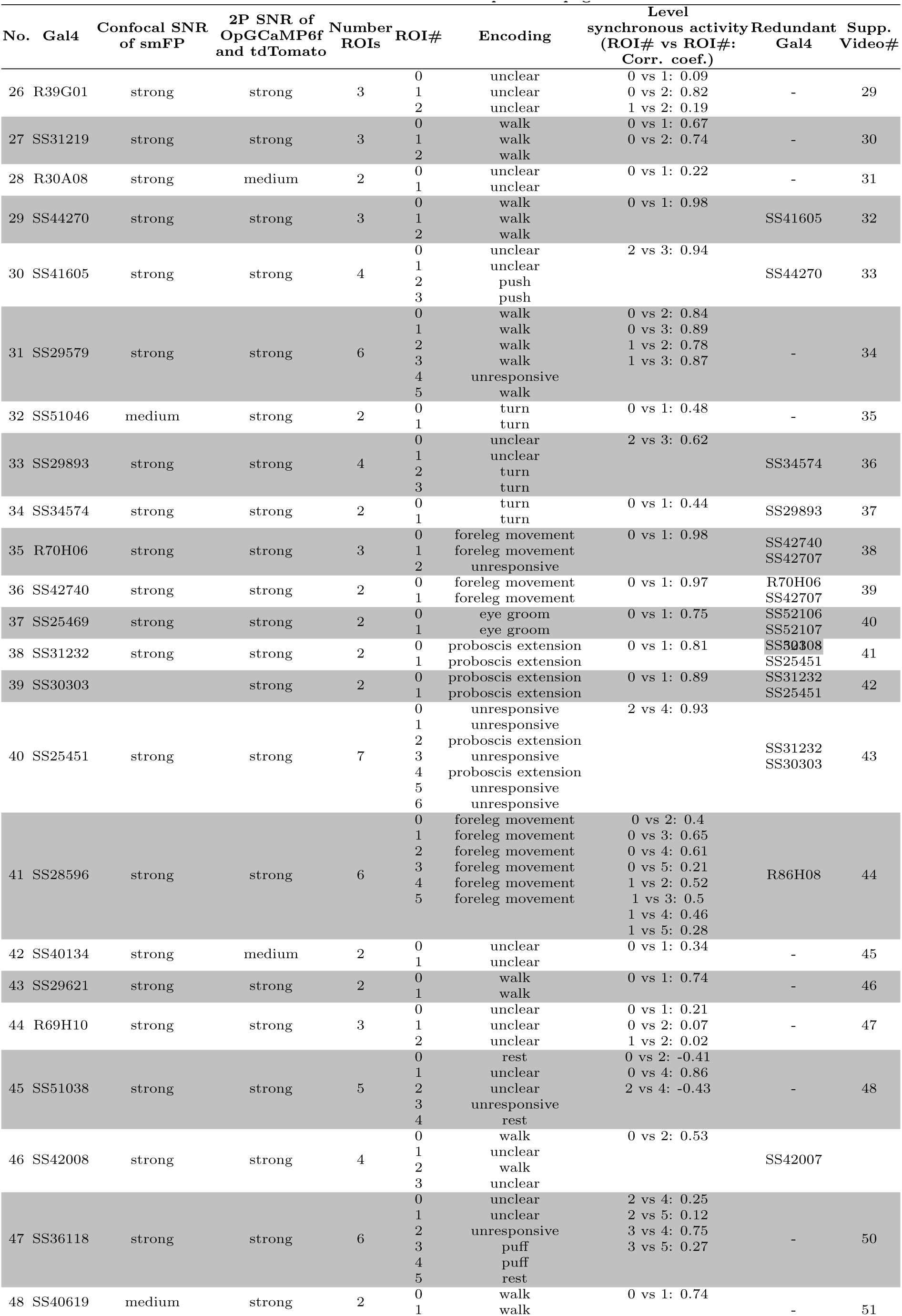
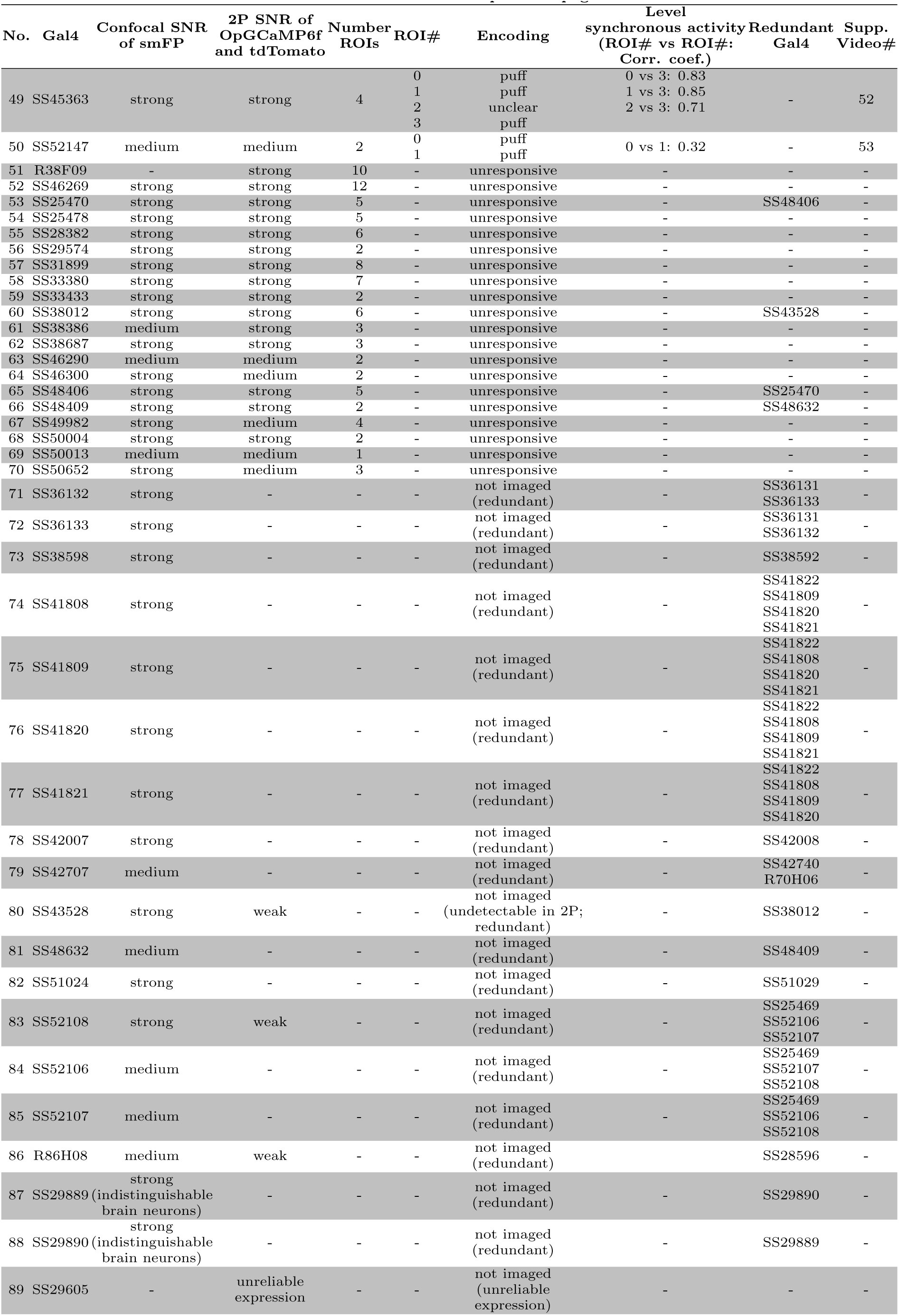
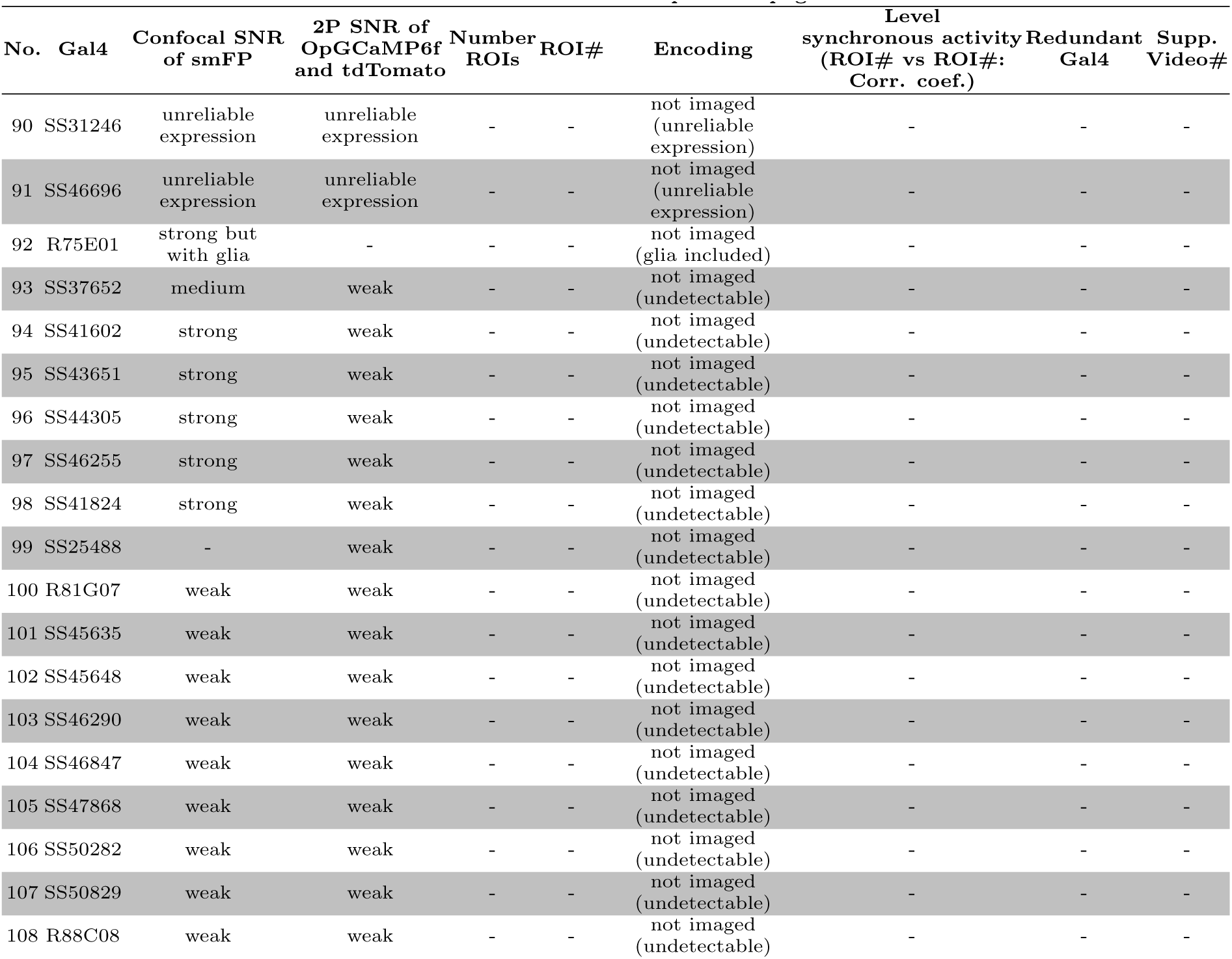
Sparse AN driver lines and associated properties.

These analyses uncovered a number of fundamental characteristics of ANs. First, as a population, ANs do not project broadly across the brain but principally target two hubs: (i) the anterior ventrolateral protocerebrum (AVLP), a site for higher-order multimodal convergence—vision [30], olfaction [31], audition [32–34], and taste [35]—, and (ii) the gnathal ganglion (GNG), a region important for action selection [23, 36, 37]. Second, ANs encode high-level behavioral states, primarily walking, rather than low-level joint or limb movements. Third, distinct behavioral state signals are system-atically conveyed to different brain targets. The AVLP is informed of self-motion states like resting, walking, and the presence of gust-like stimuli, likely to contextualize sensory cues. By contrast, the GNG receives precise signals about actions—turning, eye grooming, and proboscis extension—likely to guide action selection.

To understand the relationship between AN behavioral state encoding and brain projection patterns, we then performed a more in-depth investigation of seven AN classes. We observed a correspondence between the morphology of ANs in the VNC and their behavioral state encoding: ANs with neurites targeting all three VNC neuromeres (T1-T3) encode global locomotor states (e.g., resting and walking) while those with projections only to the T1 prothoracic neuromere encoded foreleg-dependent behaviors (e.g., eye grooming). Notably, AN axons were also present within the VNC. This suggests that ANs are not simply passive relays of behavioral state signals to the brain but that they may also help to orchestrate motor actions and/or compute state encodings. This latter possibility is illustrated by a class of ‘PE-ANs’ that seems to encode the number of proboscis extensions generated over tens of seconds, possibly through recurrent interconnectivity within the VNC. In summary, these data provide a first comprehensive view of ascending signals to the brain, opening the door for a cellular-level understanding of how behavioral states are computed, and how ascending motor signals enable the brain to contextualize sensory signals and select appropriate future actions.

## 2 Results

### 2.1 A large-scale screen of ascending neuron movement encoding, brain targeting, and motor system patterning

We performed a functional screen of 108 driver lines that target small sets of ANs (**Figure 1B**) to address to what extent they encode low-level joint and limb movements, or high-level behavioral states. To quantify limb movements, we recorded each fly using six synchronized cameras (a seventh camera was used to position the fly on the ball) (**Figure 1D**). We processed these videos using DeepFly3D [28], a markerless 3D pose estimation software that outputs joint positions and angles (**Figure 1E**). We also measured spherical treadmill rotations using two optic flow sensors [29] and converted these into three fly-centric velocities: forward (mm/s), sideways (mm/s), and yaw (degree/s) (**Figure 1F**) that correspond to forward/backward walking, side-slip, and turning, respectively. A separate DeepLabCut [38] deep neural network was used to track proboscis extensions (PEs) from one camera view **(Figure S2)**. We used a puff of CO_2_ to elicit behavior in sedentary animals.

Synchronized with movement quantification, we recorded the activity of ANs by performing two-photon imaging of the cervical connective within the thoracic ventral nerve cord (VNC) [27]. The VNC houses motor circuits that are functionally equivalent to those in the vertebrate spinal cord **(Figure 1G, left)**. Neural activity was read-out as changes in the fluorescence of a genetically-encoded calcium indicator, OpGCaMP6f, expressed in a small number of ANs. Simultaneously, we recorded tdTomato fluorescence as an anatomical fiduciary. Imaging coronal (x-z) sections of the cervical connective allowed us to keep AN axons within the imaging field-of-view despite behaviorally-induced motion artifacts that would disrupt conventional horizontal (x-y) section imaging [27]. Sparse sp-Gal4 and Gal4 fluorescent reporter expression facilitated axonal region-of-interest (ROI) detection. To semi-automatically segment and track AN ROIs across thousands of imaging frames, we developed and used AxoID, a deep network-based software **(Figure 1G, right)**(see Methods). AxoID also helped perform ROI detection despite significant movement-related ROI translations and deformations as well as, for some driver lines, relatively low transgene expression levels and suboptimal imaging signal-to-noise ratios (SNR).

To relate AN neural activity with ongoing limb movements, we trained classifiers using 3D joint angles and spherical treadmill rotational velocities to accurately and automatically detect nine behaviors—forward and backward walking, spherical treadmill pushing, resting, eye and antennal grooming, foreleg and hindleg rubbing, and abdominal grooming (**Figure 1H**). Additionally, we classified non-orthogonal, co-occurring behaviors like proboscis extensions (PEs) and recorded the timing of CO_2_ puff stimuli **(Video 1)**.

Our final dataset consisted of neural activity recordings from 247 ANs targeted using 70 sparsely-labelled driver lines (more than 32 h of data). These data included (i) anatomical projection patterns, and temporally synchronized (ii) neural activity, (iii) joint angles, and (iv) spherical treadmill rotations. Here we focus on the results for 157 of the most active ANs taken from 50 driver lines (more than 23 h of data) **(Video 2)**. The remainder were excluded due to redundancy with other driver lines, a lack of neural activity, or a low SNR (as determined by smFP confocal imaging, or two-photon imaging of tdTomato and OpGCaMP6f). Representative data from each of these selected driver lines illustrate the richness of our dataset **(Videos 3-52)**.

### 2.2 Ascending neurons encode high-level behaviors

With these data, we first asked to what extent AN activity encode low-level joint angles and leg movements, or high-level behaviors like walking, resting, and grooming **(Figure S1A)**. We expected that, unlike primary limb mechanosensory neurons, second- and higher-order ANs would more likely integrate and process proprioceptive and tactile sensory signals to encode high-level behavioral states. This remained unknown because previous studies of AN encoding [21–23] did not quantify movements at high enough resolution, or study more than a few ANs in total. To address this gap, with the data from our large-scale functional screen, we performed a linear regression analysis to quantify the degree to which the movements of individual joints, legs, pairs of legs, or epochs of high-level behaviors could explain the time-course of AN activity. Specifically, we quantified the unique explained variance (UEV, or Δ*R*^2^) for each movement, or behavioral regressor via cross-validation by subtracting a reduced model *R*^2^ from the full model *R*^2^. In the reduced model, a regressor of interest was shuffled while keeping the other regressors intact (see Methods). To compensate for the temporal mismatch between fast leg movements and slower calcium signal decay dynamics, every joint angle and behavioral state regressor was convolved with a decay kernel chosen to maximize the explained variance in neural activity.

First we examined to what extent individual joint angles could explain the activities of 157 ANs. We confirmed that the vast majority of joint angles do not covary with others—with the exception of the middle and hindleg CTr and FTi pitch angles which were highly correlated to one another **(Figure S3)**. This is important because if two regressors are highly correlated, one regressor can compensate when shuffling the other, resulting in a false negative outcome. We did not find any evidence of joint angles explaining AN activity (**Figure 2A**). Similarly, individual leg movements (tested by shuffling all of the joint angle regressors for a given leg) could not explain the variance of AN activity (**Figure 2B**). Additionally, with the exception of ANs from SS25469 whose activities could be explained by movements of the forelegs (**Figure 2C**), AN activity largely could not be explained by the movements of pairs of legs. By contrast, the activity of ANs could be explained by high-level behavioral states (**Figure 2D**). Most ANs encoded self-motion—forward walking and resting—but some also encoded specific actions like eye grooming, proboscis extensions, as well as responses to puff stimuli.

**Figure 2:**
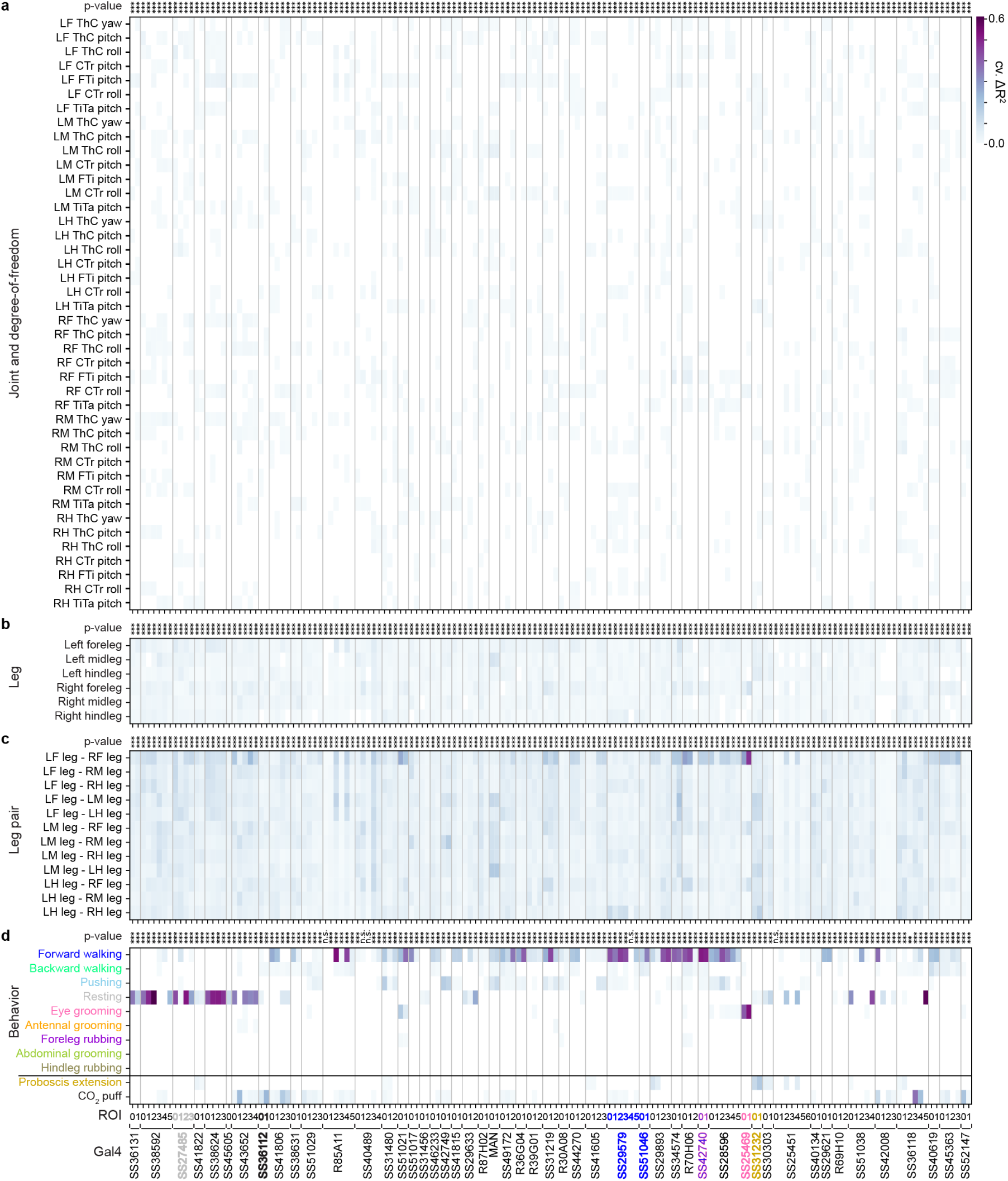
Ascending neurons encode high-level behaviors. Proportion of variance in AN activity that is uniquely explained by regressors (cross-validated Δ*R*^2^) based on **(a)** joint movements, **(b)** the movements of individual legs, **(c)** the movements of pairs of legs, **(d)** high-level behaviors. Regression analyses were performed for 157 ANs recorded from 50 driver lines. Lines selected for more in-depth analysis are color-coded by the behavioral class best explaining their neural activity: SS27485 (resting), SS36112 (puff responses), SS29579 (walking), SS51046 (turning), SS42740 (foreleg movements), SS25469 (eye grooming), and SS31232 (proboscis extensions). Non-orthogonal regressors (PE and CO_2_ puffs) are separated from the others. *P* -values report the F-statistic of overall significance of the complete regression model with none of the regressors shuffled (\**p<*0.05, \*\**p<*0.01, and \*\*\**p<*0.001)

Our regression approach is conservative and avoids false positives. However, because is prone to false negatives for infrequently occurring behaviors like abdominal grooming and hindleg rubbing, as an additional alternative approach, we measured the mean normalized Δ*F/F* for each AN for each high-level behavioral state. Using this complementary approach, we could confirm and extend our results **(Figure S4)**. We considered results from both our linear regression and mean normalized Δ*F/F* analyses when selecting neurons for further in-depth analyses.

### 2.3 Ascending neurons target integrative sensory, or action selection brain regions as a function of their encoding

Having identified high-level behavioral state encoding for a large population of ANs, we next wondered to what extent these distinct state signals are routed to specific and distinct brain targets **(Figure S1B)**. On one hand, individual ANs might project diffusely to multiple brain regions. Alternatively, they might target one, or only a few regions. For instance, locomotor signals carried by walking and resting encoding ANs might be conveyed to brain regions to contextualize time-varying visual and olfactory cues with respect to an animal’s own self-motion. On the other hand, ANs that signal when an animal is grooming might target action selection brain regions to prohibit future actions that might result in unstable postures. To address these possibilities, we quantified the brain projections of all 157 ANs by staining and imaging the expression of spFP and MCFO reporters in these neurons (**Figure 1C**).

Strikingly, we found that AN projections to the brain were largely restricted to two regions: the AVLP, a site known for multimodal, integrative sensory processing [30–35] and the GNG, a hub for action selection [23, 36, 37] (**Figure 3A**). ANs encoding resting and puff-responses almost exclusively target the AVLP **(Figure S5A,B)** providing a robust means for interpreting whether sensory cues arise from self-motion or the movement of objects in the external environment: while resting, an animal can perceive visual motion due to moving objects, and odor fluctuations due to gust-like puffs of air. By contrast, the GNG is targeted by ANs encoding a wide variety of behavioral states including walking, eye grooming, and proboscis extensions **(Figure S5A,B)**. These signals may ensure that future actions are compatible with ongoing ones.

**Figure 3:**
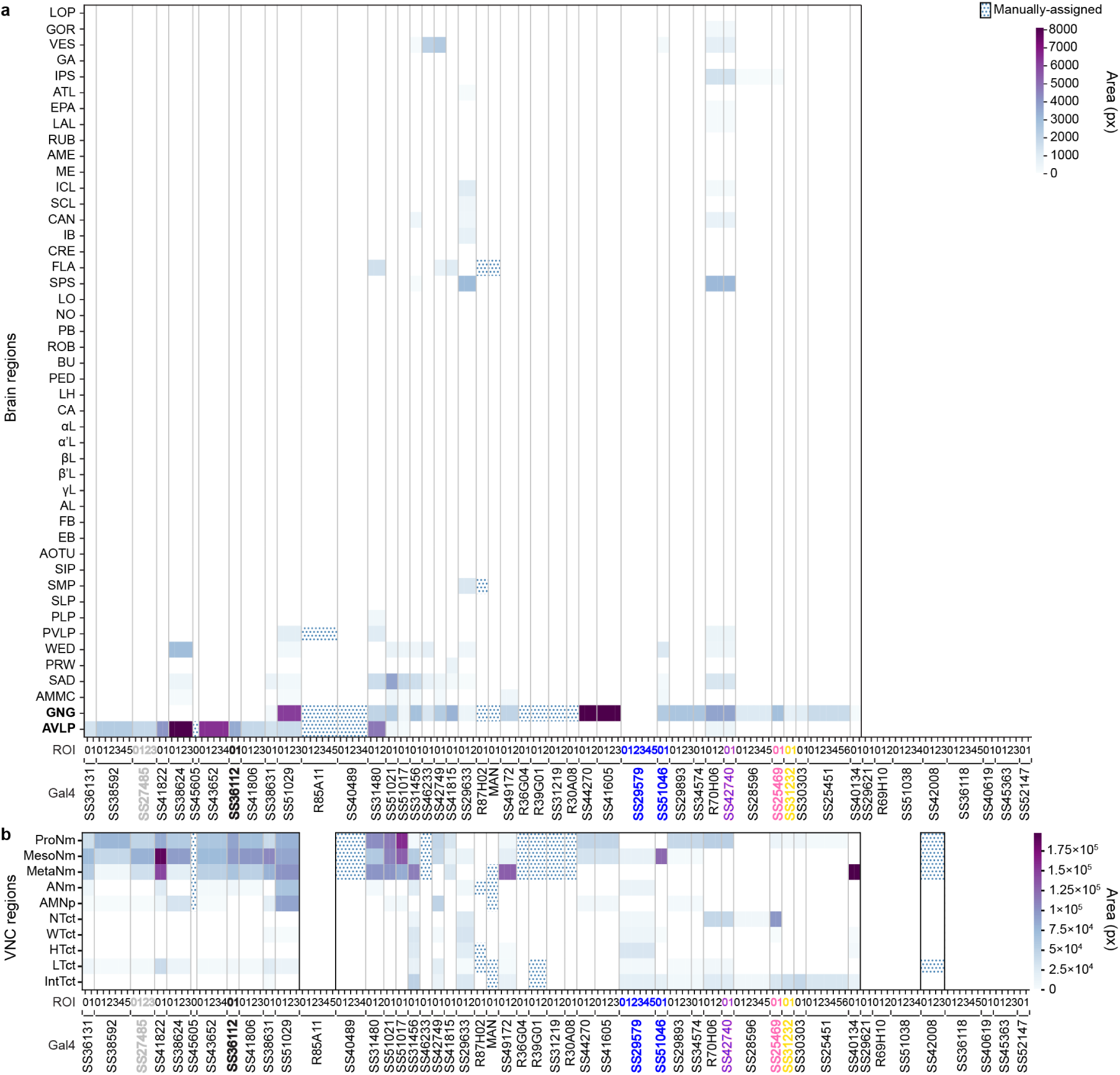
Ascending neurons principally project to the brain’s AVLP and GNG and the VNC’s leg neuromeres. Regional innervation of **(a)** the brain, or **(b)** the VNC. Data are for 157 ANs recorded from 50 driver lines and quantified through pixel-based analyses of MCFO labeled confocal images. Manually quantified driver lines are indicated (dotted). Lines for which projections could not be unambiguously identified are left blank. Lines selected for more in-depth evaluation are color-coded by the behavioral state that best explains their neural activity: SS27485 (resting), SS36112 (puff responses), SS29579 (walking), SS51046 (turning), SS42740 (foreleg-dependent behaviors), SS25469 (eye grooming), and SS31232 (proboscis extensions).

Because AN dendrites and axons within the VNC might help to compute behavioral state signals, we next asked to what extent their projection patterns within the VNC are predictive of an AN’s encoding. For example, ANs encoding resting might require sampling each VNC leg neuromere (T1, T2, and T3) to confirm that all legs are inactive. By quantifying AN projections within the VNC (**Figure 3B**), we found that, indeed, an AN’s VNC projection pattern can be predictive of behavioral state encoding. As hypothesized, ANs encoding resting (e.g., SS27485) all project to every VNC leg neuromere **(Figure S5A,C)**. By contrast, ANs encoding foreleg-dependent eye grooming (SS25469) only project within T1, the VNC neuromere that houses motor circuits that control the front legs **(Figure S5A,C)**. Next, to more precisely investigate how the morphological features of ANs relate to behavioral state encoding, we performed a more detailed study of a diverse subset of ANs that encode resting, puff-responses, walking, turning, foreleg-dependent behaviors, eye grooming, and proboscis extensions.

### 2.4 Distinct rest- and puff-encoding by morphologically similar ANs

AN classes that encode resting and puff responses had coarsely similar projection patterns: both almost exclusively target the brain’s AVLP while also both sampling from all three VNC leg neuromeres (T1-T3) **(Figure S5)**. We therefore next investigated which more detailed morphological features might be predictive of their very divergent encoding.

We addressed this question by closely examining the functional and morphological properties of specific pairs of ‘rest-ANs’ (SS27485) and ‘puff-ANs’ (SS36112). Neural activity traces of rest-ANs and puff-ANs could be reliably predicted by regressors for resting (**Figure 4A**), and puff-stimuli (**Figure 5A**), respectively. This was statistically confirmed by comparing behavior-triggered averages of AN responses at the onset of resting (**Figure 4B**), or puff stimulation (**Figure 5B**), respectively. Importantly, although CO_2_ puffs frequently elicited brief periods of backward walking, close analysis revealed that puff-ANs primarily respond to gust-like puffs and do not encode backward walking **(Figure S6)**. They also did not encode responses to CO_2_ specifically: the same neurons responded equally well to air puffs **(Figure S7)**.

**Figure 4:**
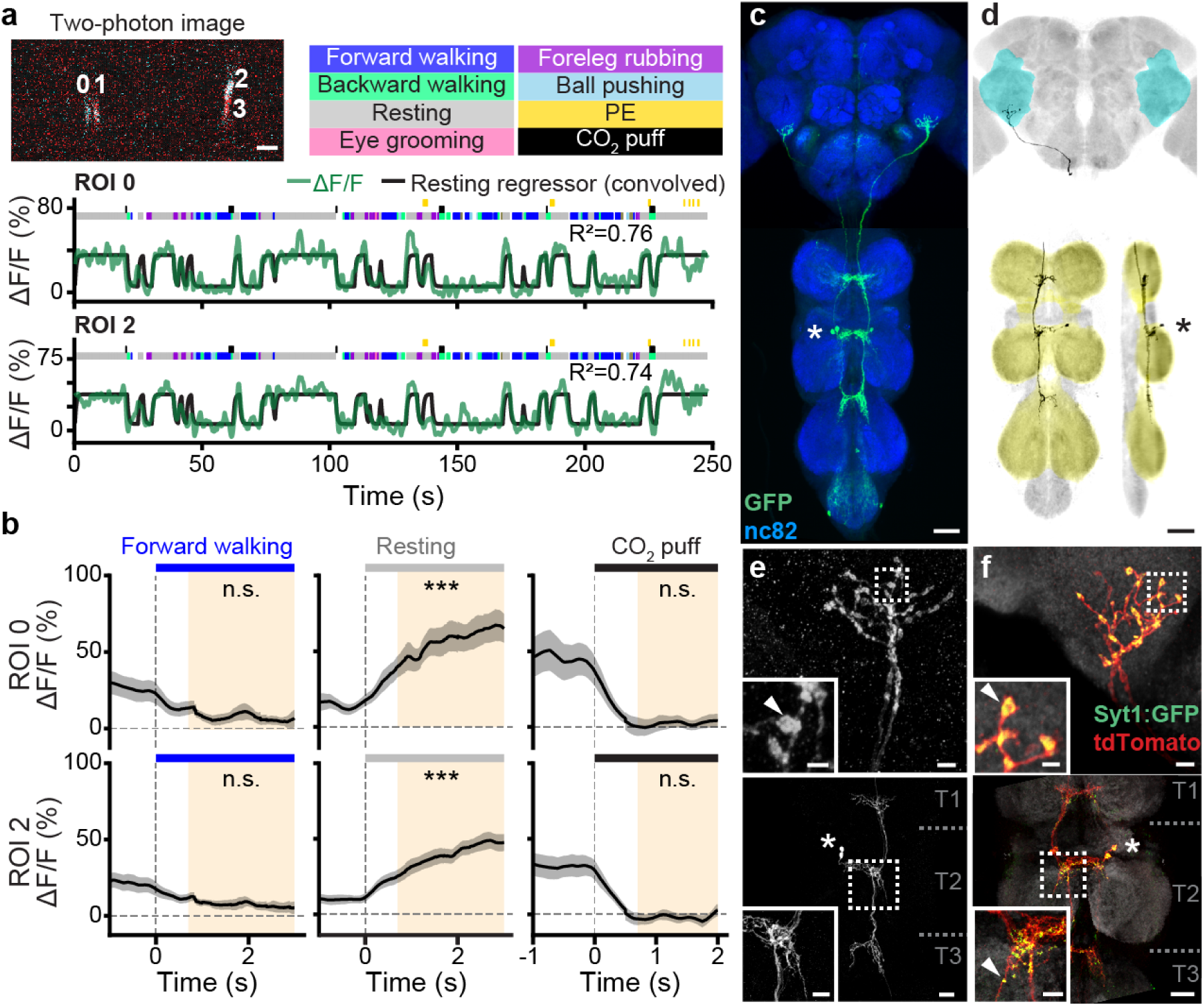
Functional and anatomical properties of ascending neurons encoding resting. **(a) (top-left)** Two-photon image of axons from an SS27485-Gal4 animal expressing OpGCaMP6f (cyan) and tdTomato (red). ROIs are numbered. Scale bar is 5 µm. **(bottom)** Behavioral epochs are color-coded. Representative Δ-F/F time-series from two ROIs (green) overlaid with a prediction (black) obtained by convolving resting epochs with a Ca^2+^ response function. Explained variance is indicated (*R*^2^). **(b)** Mean (solid line) and 95% confidence interval (gray shading) of Δ-F/F traces during epochs of forward walking (left), resting (middle), or CO_2_ puffs (right). 0 s indicates the start of each epoch. Here and in Figures 5 - 9, data more than 0.7s after onset (yellow region) are compared with an otsu thresholded baseline (ANOVA and Tukey posthoc comparison, \*\*\**p<*0.001, \*\**p<*0.01, \**p<*0.05, n.s. not significant). **(c)** Standard deviation projection image of an SS27485-Gal4 nervous system expressing smFP and stained for GFP (green) and Nc82 (blue). Cell bodies are indicated (white asterisk). Scale bar is 40 µm. **(d)** Projection as in **c** but for one MCFO-expressing, traced neuron (black asterisk). The brain’s AVLP (cyan) and VNC’s leg neuromeres (yellow) are color-coded. Scale bar is 40 µm. **(e, f)** Higher magnification projections of **(top)** brains and **(bottom)** VNCs of SS27485-Gal4 animals expressing **(e)** the stochastic label MCFO, or **(f)** the synaptic marker, syt:GFP (green), and tdTomato (red). Insets magnify dashed boxes. Indicated are cell bodies (asterisks), bouton-like structures (white arrowheads), and VNC leg neuromeres (‘T1, T2, T3’). Scale bars for brain images and insets are 5 µm and 2 µm, respectively. Scale bars for VNC images and insets are 20 µm and 10 µm, respectively.

**Figure 5:**
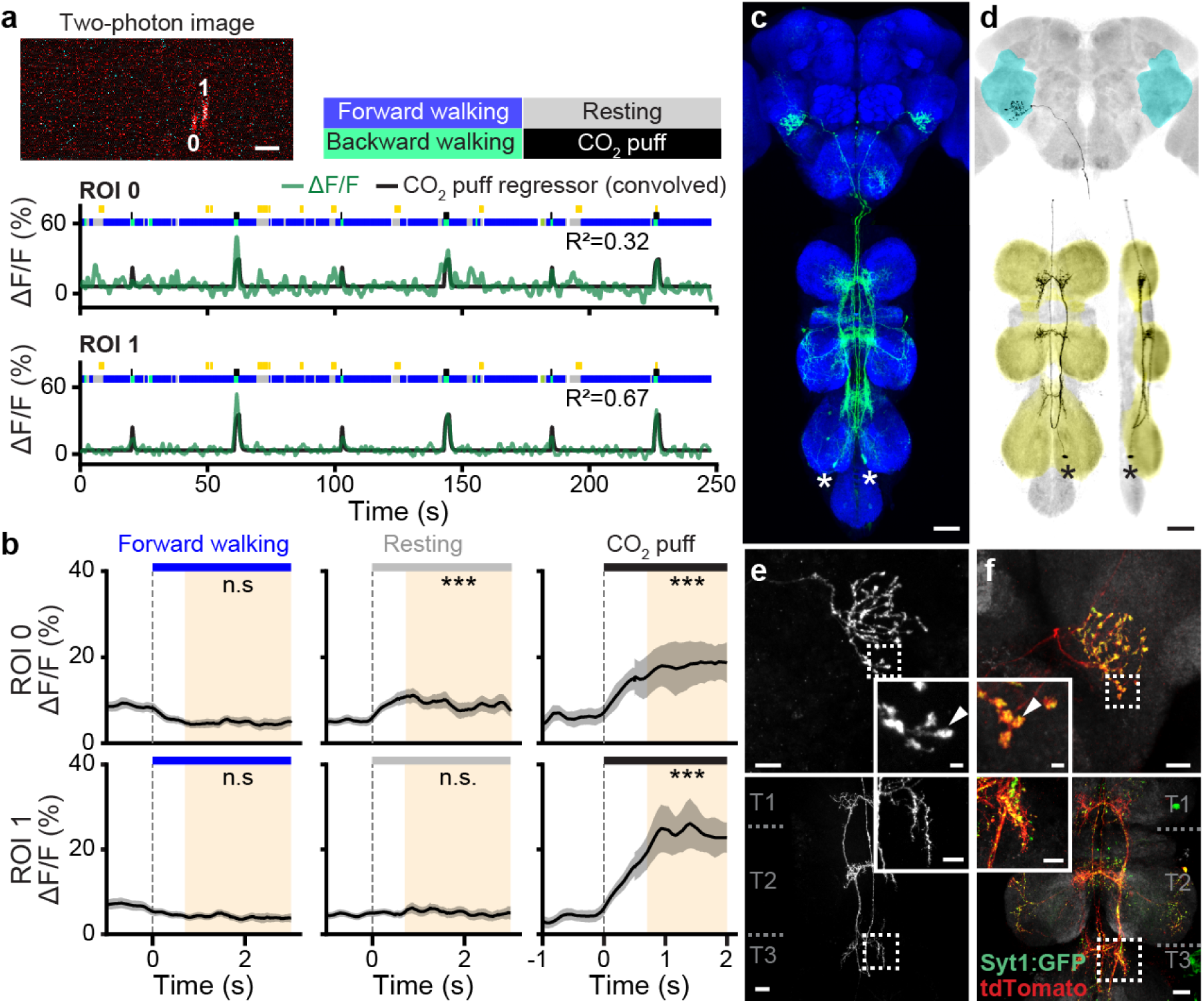
Functional and anatomical properties of ascending neurons responding to puffs. **(a) (top-left)** Two-photon image of axons from an SS36112-Gal4 animal expressing OpGCaMP6f (cyan) and tdTomato (red). ROIs are numbered. Scale bar is 5 µm. **(bottom)** Behavioral epochs are color-coded. Representative Δ-F/F time-series from two ROIs (green) overlaid with a prediction (black) obtained by convolving CO_2_ puff periods with a Ca^2+^ response function. Explained variance is indicated (*R*^2^). **(b)** Mean (solid line) and 95% confidence interval (gray shading) of Δ-F/F traces during epochs of forward walking (left), resting (middle), or CO_2_ puffs (right). 0 s indicates the start of each epoch. **(c)** Standard deviation projection image for an SS36112-Gal4 nervous system expressing smFP and stained for GFP (green) and Nc82 (blue). Cell bodies are indicated (white asterisks). Scale bar is 40 µm. **(d)** Projection as in **c** but for one MCFO-expressing, traced neuron (black asterisks). The brain’s AVLP (cyan) and VNC’s leg neuromeres (yellow) are color-coded. Scale bar is 40 µm. **(e, f)** Higher magnification projections of **(top)** brains and **(bottom)** VNCs of SS36112-Gal4 animals expressing **(e)** the stochastic label MCFO, or **(f)** the synaptic marker, syt:GFP (green), and tdTomato (red). Insets magnify dashed boxes. Indicated are bouton-like structures (white arrowheads), and VNC leg neuromeres (‘T1, T2, T3’). Scale bars for brain images and insets are 10 µm and 2 µm, respectively. Scale bars for VNC images and insets are 20 µm and 10 µm, respectively.

As mentioned, rest- and puff-ANs, despite their very distinct encoding, exhibit similar innervation patterns in the brain and VNC. However, MCFO-based single neuron analysis revealed a few subtle but important differences. First, rest- and puff-AN cell bodies are located in the T2 (**Figure 4C**) and T3 (**Figure 5C**) neuromeres, respectively. Second, although both AN classes project medially into all three leg neuromeres (T1-T3), rest-ANs have a simpler morphology (**Figure 4D**) compared with the more complex arborization of puff-ANs in the VNC (**Figure 5D**). In the brain, both AN types project to nearly the same ventral region of the AVLP. There, they exhibit varicose terminals (**Figure 4E** and **Figure 5E**). Using syt:GFP, a GFP tagged synaptotagmin (presynaptic) marker, we confirmed that these varicosities house synaptic terminals **(Figure 4F, top** and **Figure 5F, top)**. Notably, in addition to smooth, likely dendritic arbors, both AN classes have axon terminals within the VNC **(Figure 4F, bottom** and **Figure 5F, bottom)**.

Taken together, these results demonstrate that even very subtle differences in VNC patterning can give rise to dramatically different AN tuning properties. In the case of rest- and puff-ANs, we speculate that this might be due to physically close, but distinct presynaptic partners—possibly leg proprioceptive afferents for rest-ANs, and leg tactile afferents for puff-ANs.

### 2.5 Walk- or turn-encoding depends on the laterality of VNC projections

Among the ANs we analyzed, most encoded walking (**Figure 2D**). However, this broad category of locomotion includes more subtle dimensions including walking direction and turning. We reasoned that an AN’s patterning within the VNC may be predictive of whether it encodes locomotion broadly (e.g., walking) versus narrowly (e.g., turning).

Indeed, we observed that while the activity of one pair of ANs (SS29579, ‘walk-ANs’) was remarkably well explained by the timing and onset of walking epochs (**Figure 6A-C**), for other ANs a broad walking regressor could account for much less variance in neural activity (**Figure 2D**). We reasoned that these ANs might instead encode narrower locomotor features like turning. For example, for a bilateral pair of DNa01 descending neurons, their difference in activity correlates with turn direction [27, 39]. To see if this might also be the case for some pairs of walk-encoding ANs, we quantified the degree to which their difference in activity can be explained by spherical treadmill roll and yaw velocities—a proxy for turning behaviors (**Figure 7A**). Indeed, we found one pair of ANs (SS51046) for which turning explained a relatively large amount of variance. For this pair of ‘turn-ANs’, although a combination of forward and backward walking regressors poorly predicted neural activity (**Figure 7B**), a regressor based on spherical treadmill roll velocities strongly predicted the difference in activity between this bilateral pair of neurons (**Figure 7C**). When an animal turned right, the right (ipsilateral) turn-AN was active. Conversely, the left turn-AN was active during left turns (**Figure 7D**). During forward walking, both turn-ANs were active (**Figure 7E**).

**Figure 6:**
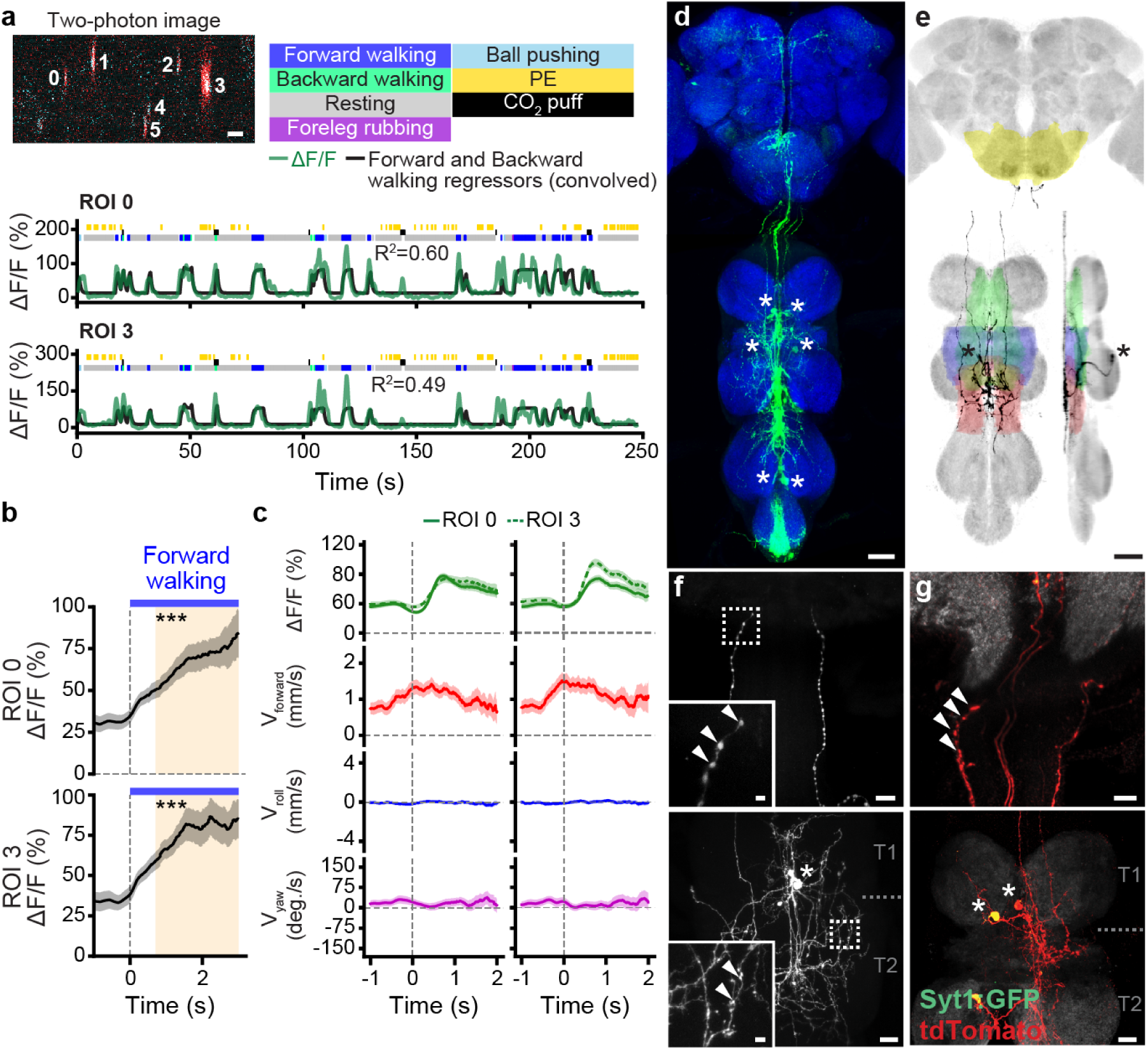
Functional and anatomical properties of ascending neurons encoding walking. **(a) (top-left)** Two-photon image of axons from an SS29579-Gal4 animal expressing OpGCaMP6f (cyan) and tdTomato (red). ROIs are numbered. Scale bar is 5 µm. **(bottom)** Behavioral epochs are color-coded. Representative Δ-F/F time-series from two ROIs (green) overlaid with a prediction (black) obtained by convolving forward walking epochs with a Ca^2+^ response function. Explained variance is indicated (*R*^2^). **(b)** Mean (solid line) and 95% confidence interval (gray shading) of Δ-F/F traces during epochs of forward walking. 0 s indicates the start of each epoch. **(c)** Fluorescence (OpGCaMP6f) event-based ball rotations for **(left)** ROI 3, or **(right)** ROI 0. Fluorescence events are time-locked to 0 s (green). Shown are mean and 95% confidence intervals for forward (red), roll (blue), and yaw (purple) ball rotational velocities. **(d)** Standard deviation projection image for a SS29579-Gal4 nervous system expressing smFP and stained for GFP (green) and Nc82 (blue). Cell bodies are indicated (white asterisks). Scale bar is 40 µm. **(e)** Projection as in **d** but for one MCFO-expressing, traced neuron (black asterisks). The brain’s GNG (yellow) and VNC’s intermediate (green), wing (blue), and haltere (red) tectulum are color-coded. Scale bar is 40 µm. **(f, g)** Higher magnification projections of **(top)** brains and **(bottom)** VNCs of SS29579-Gal4 animals expressing **(f)** the stochastic label MCFO, or **(g)** the synaptic marker, syt:GFP (green), and tdTomato (red). Insets magnify dashed boxes. Indicated are cell bodies (asterisks), bouton-like structures (white arrowheads), and VNC leg neuromeres (‘T1, T2’). Scale bars for brain images and insets are 10 µm and 2 µm, respectively. Scale bars for VNC images and insets are 20 µm and 4 µm, respectively.

**Figure 7:**
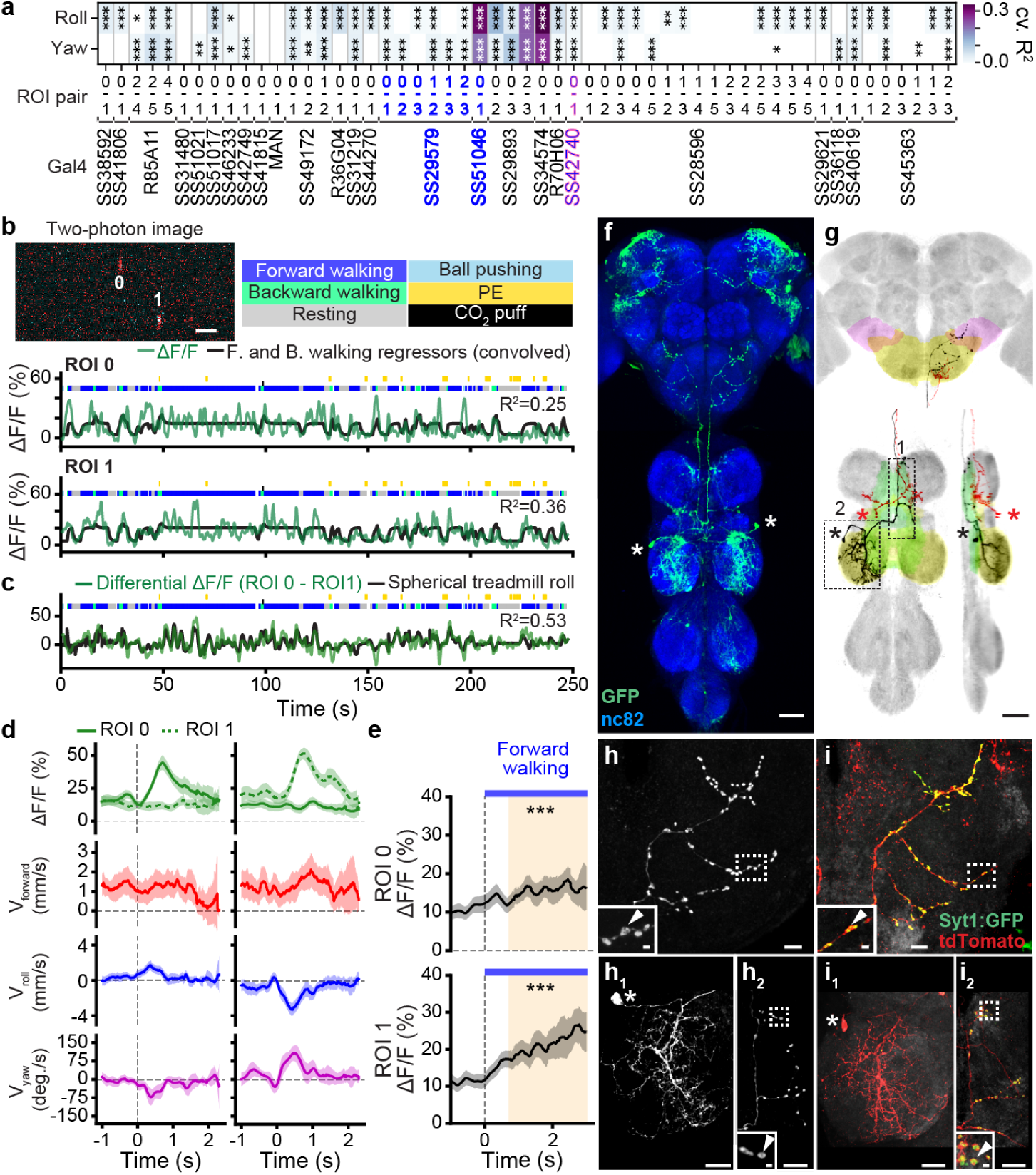
Functional and anatomical properties of ascending neurons encoding turning. **(a)** Variance explained by side-slip and turning for driver lines encoding forward walking. **(b) (top-left)** Two-photon image of axons from an SS51046-Gal4 animal expressing OpGCaMP6f (cyan) and tdTomato (red). ROIs are numbered. Scale bar is 5 µm. **(bottom)** Behavioral epochs are color-coded. Representative Δ-F/F time-series from two ROIs (green) overlaid with a prediction (black) obtained by convolving forward walking epochs with a Ca^2+^ response function. Explained variance is indicated (*R*^2^). **(c)** The differential Δ-F/F time-series obtained by subtracting the two ROIs (green) is overlaid with a prediction (black) from spherical treadmill roll rotations convolved with a Ca^2+^ response function. Explained variance is indicated (*R*^2^). **(d)** Fluorescence (OpGCaMP6f) event-based ball rotations for **(left)** ROI 0, or **(right)** ROI 1. Fluorescence events are time-locked to 0 s (green). Shown are mean and 95% confidence intervals for forward (red), roll (blue), and yaw (purple) ball rotational velocities. **(e)** Mean (solid line) and 95% confidence interval (gray shading) of Δ-F/F traces during epochs of forward walking. 0 s indicates the start of each epoch. **(f)** Standard deviation projection image for a SS51046-Gal4 nervous system expressing smFP and stained for GFP (green) and Nc82 (blue). Cell bodies are indicated (white asterisks). Scale bar is 40 µm. **(g)** Projection as in **f** but for one MCFO-expressing, traced neuron (black asterisks). The brain’s GNG (yellow), wedge (pink), and VNC’s intermediate tectulum (green), and mesothoracic leg neuromere (yellow), are color-coded. Scale bar is 40 µm. **(h, i)** Higher magnification confocal z-projections of **(top)** brains and **(bottom)** VNCs of SS51046-Gal4 animals expressing **(h)** the stochastic label MCFO, or **(i)** the synaptic marker, syt:GFP (green), and tdTomato (red). Insets magnify dashed boxes. Indicated are cell bodies (asterisks), bouton-like structures (white arrowheads), and VNC leg neuromeres (‘T1, T2’). Scale bars for brain images and insets are 10 µm and 2 µm, respectively. Scale bars for VNC images and insets are 20 µm and 2 µm, respectively.

We next asked how VNC patterning might predict this distinction between broad (walk-AN) versus narrow (turn-AN) locomotor encoding. Both AN classes have cell bodies in the VNC’s T2 neuromere (**Figure 6D** and **Figure 7F**). However, walk-ANs bilaterally innervate the T2 neuromere (**Figure 6E**), whereas turn-ANs unilaterally innervate T1 and T2 **(Figure 7G, black)**. Their ipsilateral T2 projections are smooth and likely dendritic **(Figure 7H_1_,I_1_)**, while their contralateral T1 projections are varicose and exhibit syt:GFP puncta, suggesting that they harbor presynaptic terminals **(Figure 7H_2_,I_2_)**. Both walk-ANs (**Figure 6D,E**) and turn-ANs (**Figure 7F,G**) project to the brain’s GNG. However, only turn-ANs project to the WED (**Figure 7H,I**). Notably, walk-AN terminals in the brain (**Figure 6F**) are not labelled by syt:GFP (**Figure 6G**), suggesting that they may be neuromodulatory in nature and rely on another class of synaptic machinery [40].

These data support the notion that broad versus narrow behavioral state encoding of ANs may arise from the laterality of VNC patterning. Additionally, we observed that pairs of broadly-tuned walk-ANs that bilaterally innervate the VNC are synchronously active. By contrast, pairs of narrowly-tuned turn-ANs are asynchronously active. This correlation between the laterality of an AN pair’s VNC projections and their synchrony seems to be a general principle **(Figure S8)**.

### 2.6 Foreleg-dependent actions are encoded by ANs in the anterior VNC

In addition to locomotion, flies use their forelegs to perform complex movements including reaching, boxing, courtship tapping, and several kinds of grooming—eye grooming, antennal grooming, and foreleg rubbing. An ongoing awareness of these behavioral states is critical to select appropriate future actions that do not lead to instability. For example, before deciding to groom its hindlegs, an animal must first confirm that its forelegs are stably on the ground and not also grooming.

We noted that some ANs project only to the VNC’s anterior-most, T1 leg neuromere **(Figure S5C)**. This pattern implied a potential role in encoding actions that only depend on the forelegs. Indeed, close examination revealed two classes of ANs that encode foreleg-related behaviors. We found ANs (SS42740) that were broadly active during multiple foreleg-dependent behaviors including walking, pushing, and grooming (‘foreleg-ANs’; overlaps with R70H06) **(Figure S4)(Figure 8A,B)**. By contrast, another pair of ANs (SS25469) were narrowly and sometimes asynchronously active only during eye grooming (‘eye groom-ANs’) **(Figure S4) (Figure 9A,B)**. Similar to walking and turning, we hypothesized that this broad (foreleg) versus narrow (eye groom) behavioral encoding might be reflected by a difference in the promiscuity and laterality of AN innervations in the VNC.

**Figure 8:**
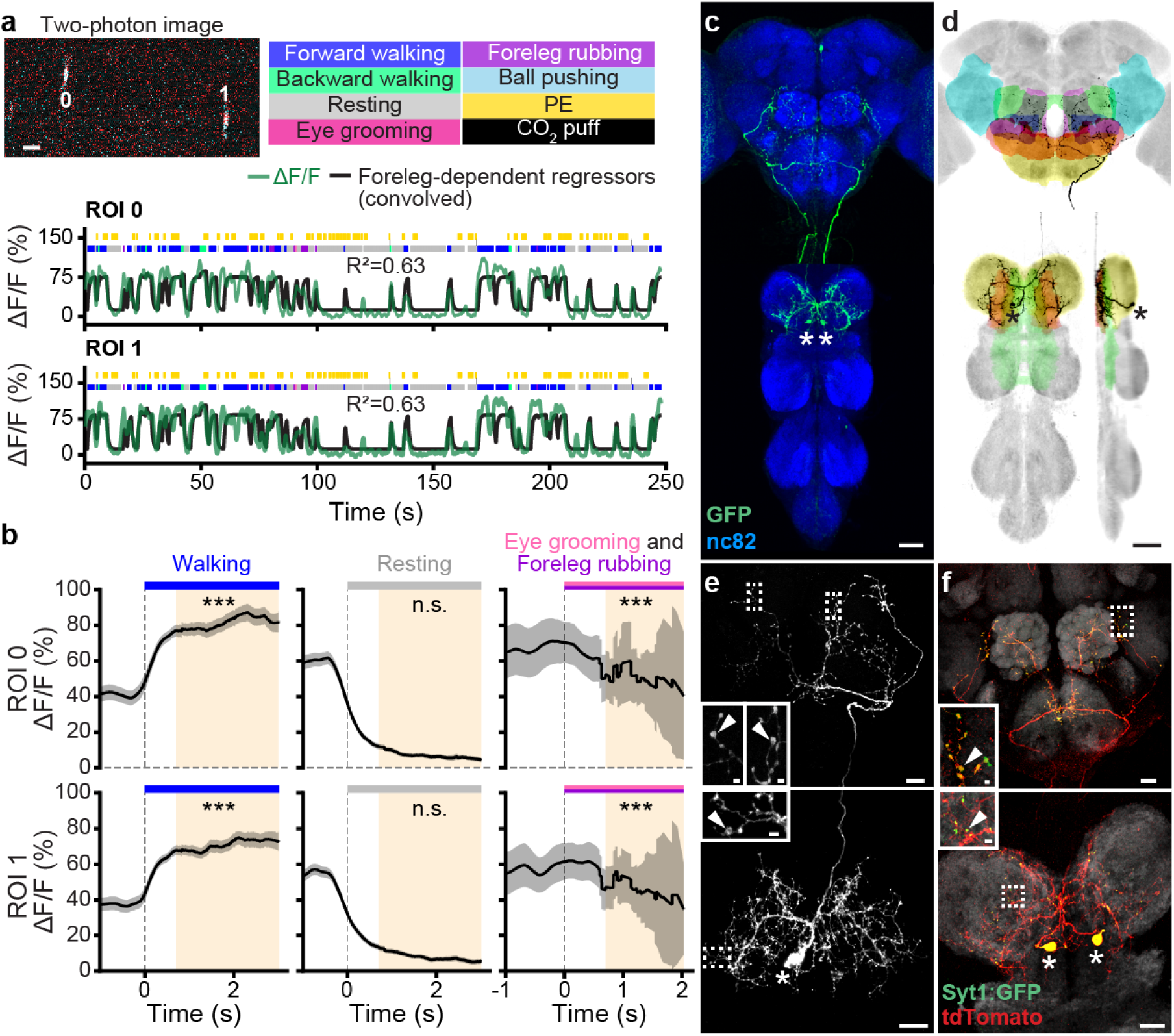
Functional and anatomical properties of ascending neurons encoding foreleg-dependent behaviors. **(a) (top-left)** Two-photon image of axons from an SS42740-Gal4 animal expressing OpGCaMP6f (cyan) and tdTomato (red). ROIs are numbered. Scale bar is 5 µm. **(bottom)** Behavioral epochs are color-coded. Representative Δ-F/F time-series from two ROIs (green) overlaid with a prediction (black) obtaind by convolving all foreleg-dependent behavioral (forward and backward walking as well as eye, antennal, and foreleg grooming) epochs with a Ca^2+^ response function. Explained variance is indicated (*R*^2^). **(b)** Mean (solid line) and 95% confidence interval (gray shading) of Δ-F/F traces during epochs of forward walking (left), resting (middle), or eye grooming and foreleg rubbing (right). 0 s indicates the start of each epoch. **(c)** Standard deviation projection image for an SS42740-Gal4 nervous system expressing smFP and stained for GFP (green) and Nc82 (blue). Cell bodies are indicated (white asterisks). Scale bar is 40 µm. **(d)** Projection as in **c** but for one MCFO-expressing, traced neuron (black asterisks). The brain’s GNG (yellow), AVLP (cyan), SAD (green), VES (pink), IPS (blue), SPS (orange), and VNC’s neck and intermediate tectulum (orange and green, respectively), and prothoracic leg neuromere (yellow) are color-coded. Scale bar is 40 µm. **(e, f)**Higher magnification confocal z-projections of **(top)** brains and **(bottom)** VNCs for SS42740-Gal4 animals expressing **(e)** the stochastic label MCFO, or **(f)** the synaptic marker, syt:GFP (green), and tdTomato (red). Insets magnify dashed boxes. Indicated are cell bodies (asterisks), and bouton-like structures (white arrowheads). Scale bars for brain images and insets are 20 µm and 2 µm, respectively. Scale bars for VNC images and insets are 20 µm and 2 µm, respectively.

**Figure 9:**
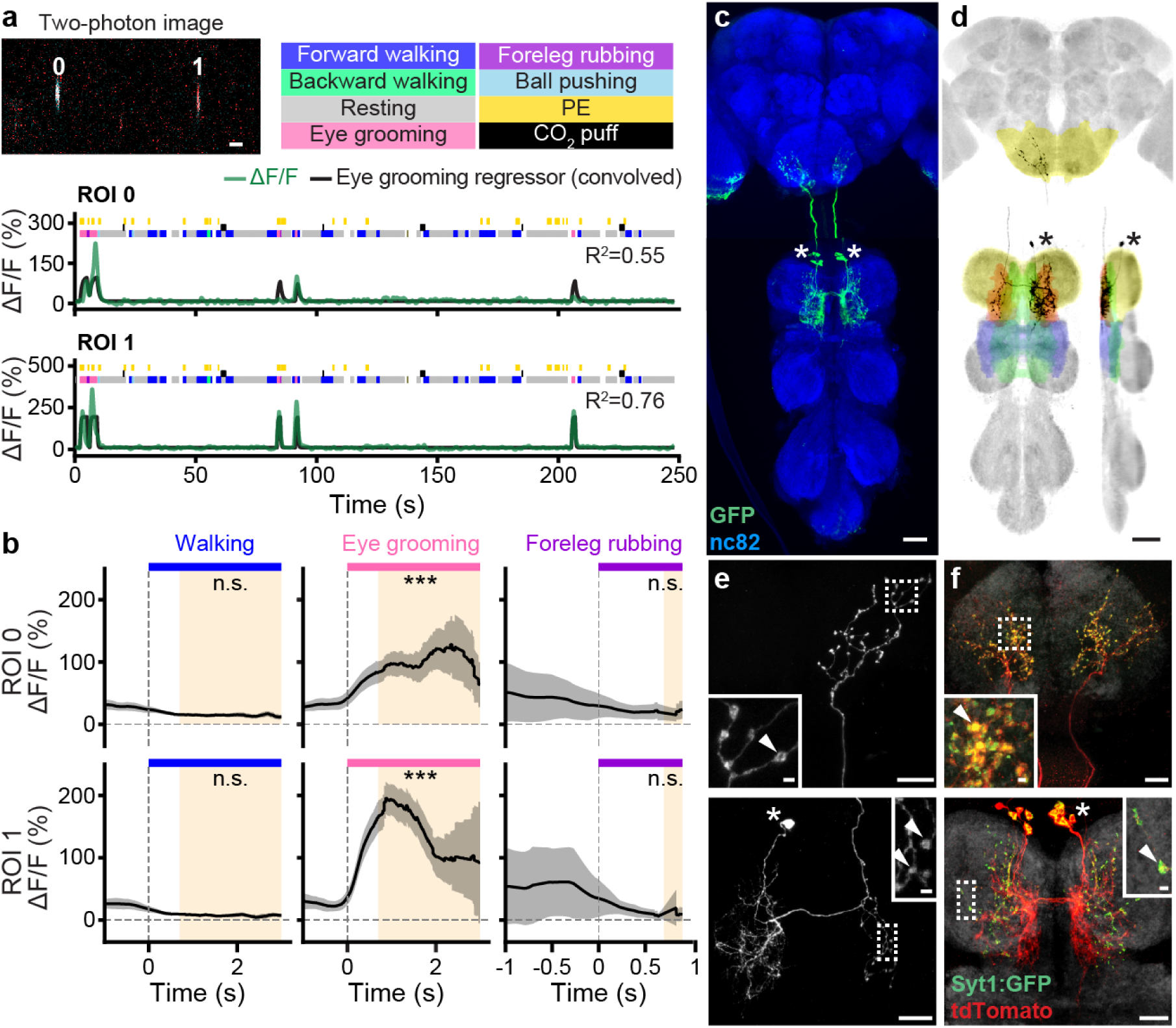
Functional and anatomical properties of ascending neurons encoding eye grooming. **(a) (top-left)** Two-photon image of axons from an SS25469-Gal4 animal expressing OpG-CaMP6f (cyan) and tdTomato (red). ROIs are numbered. Scale bar is 5 µm. **(bottom)** Behavioral epochs are color-coded. Representative Δ-F/F time-series from two ROIs (green) overlaid with a prediction (black) obtained by convolving eye grooming epochs with a Ca^2+^ response function. Explained variance is indicated (*R*^2^). **(b)** Mean (solid line) and 95% confidence interval (gray shading) of Δ-F/F traces during epochs of forward walking (left), eye grooming (middle), or foreleg rubbing (right). 0 s indicates the start of each epoch. **(c)** Standard deviation projection image for an SS25469-Gal4 nervous system expressing smFP and stained for GFP (green) and Nc82 (blue). Cell bodies are indicated (white asterisks). Scale bar is 40 µm. **(d)** Projection as in **c** but for one MCFO-expressing, traced neuron (black asterisks). The brain’s GNG (yellow) and VNC’s intermediate, neck and wing tectulum (green, red, and blue respectively), and prothoracic leg neuromere (yellow) are color-coded. Scale bar is 40 µm. **(e, f)** Higher magnification projections of **(top)** brains and **(bottom)** VNCs for SS25469-Gal4 animals expressing **(e)** the stochastic label MCFO, or **(f)** the synaptic marker, syt:GFP (green), and tdTomato (red). Insets magnify dashed boxes. Indicated are cell bodies (asterisks), and bouton-like structures (white arrowheads). Scale bars for brain images and insets are 20 µm and 2 µm, respectively. Scale bars for VNC images and insets are 20 µm and 2 µm, respectively.

To test this hypothesis, we compared the morphologies of foreleg- and eye groom-ANs. Both had cell bodies in the T1 neuromere, although foreleg-ANs were posterior (**Figure 8C**) and eye groom-ANs were anterior (**Figure 9C**). Foreleg- and eye groom-ANs also both projected to the dorsal T1 neuromere with eye groom-AN neurites restricted to the tectulum (**Figure 8D** and **Figure 9D**). Notably, foreleg-AN puncta **(Figure 8E, bottom)** and syt:GFP **(Figure 8F, bottom)** were bilateral and diffuse while eye groom-AN puncta **(Figure 9E, bottom)** and syt:GFP **(Figure 9F, bottom)** were largely restricted to the contralateral T1 neuromere. Projections to the brain paralleled this difference in VNC projection promiscuity: foreleg-ANs terminated across multiple brain areas—GNG, AVLP, SAD, VES, IPS, and SPS (**Figure 8E,F** **top)**— while eye groom-ANs narrowly targeted the GNG (**Figure 9E,F** **top)**.

These results further illustrate that AN encoding can be predicted by VNC patterning. Here, diffuse, bilateral projections are associated with encoding multiple behaviors that require foreleg movements whereas focal, unilateral projections give rise to a narrow encoding of eye grooming.

### 2.7 Temporal integration of proboscis extensions by a cluster of ANs

Flies often generate spontaneous proboscis extensions (PEs) while resting **(Figure 10A, yellow ticks)**. We observed that ‘PE-ANs’ (SS31232, overlap with SS30303) (**Figure 2D**) become active during PE trains—a sequence of PEs that occur within a short period of time (**Figure 10A**). Close examination revealed that PE-AN activity slowly ramped up over the course of PE trains. This made them difficult to model using a simple PE regressor: their activity levels were lower than predicted early in PE trains, and higher than predicted late in PE trains. On average, across many PE trains, PE-AN activity reached a plateau by the seventh PE (**Figure 10B**).

**Figure 10:**
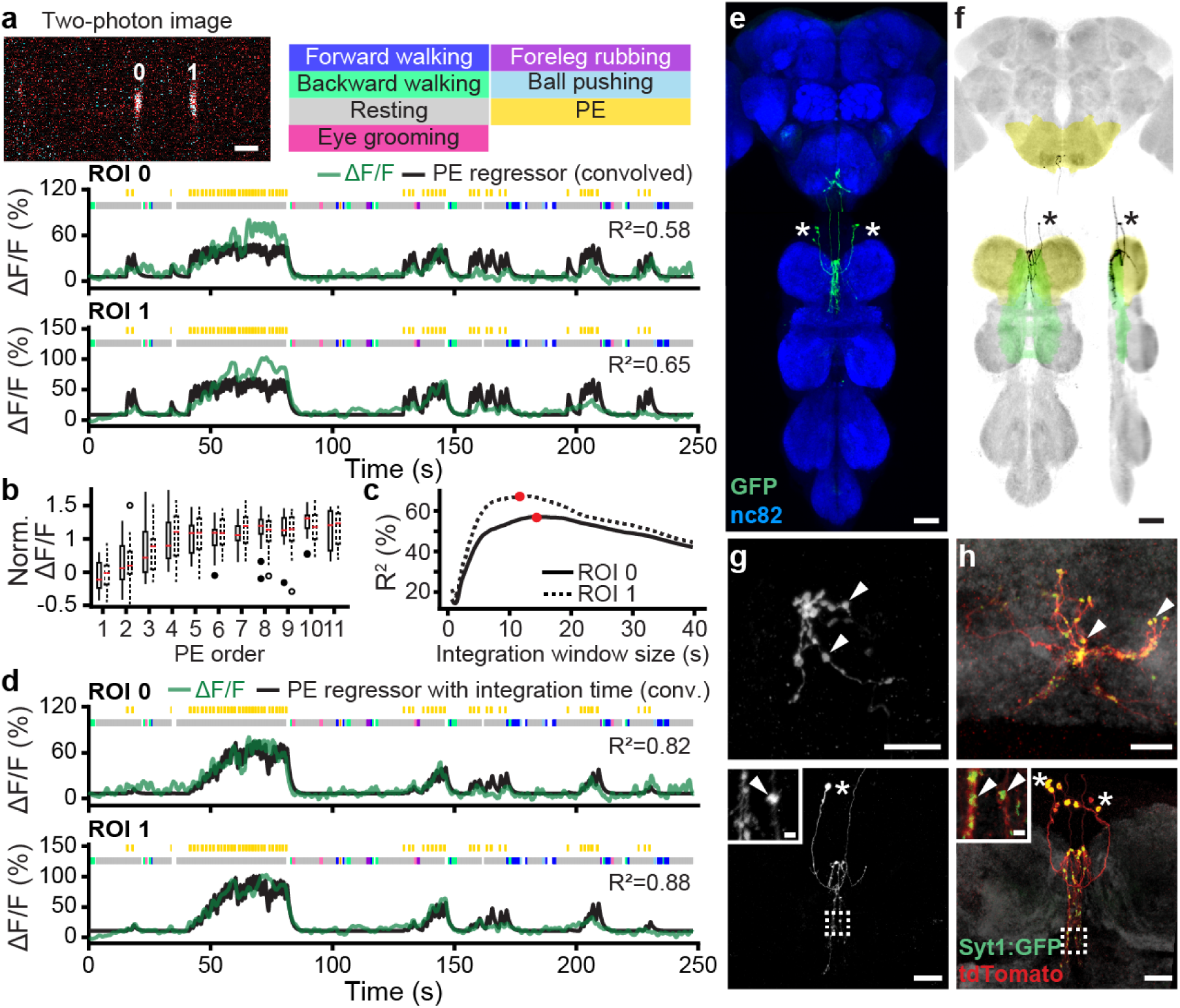
Functional and anatomical properties of ascending neurons integrating the number of proboscis extensions over time. **(a) (top-left)** Two-photon image of axons from an SS31232-Gal4 animal expressing OpGCaMP6f (cyan) and tdTomato (red). ROIs are numbered. Scale bar is 5 µm. **(bottom)** Behavioral epochs are color-coded. Representative Δ-F/F time-series from two ROIs (green) overlaid with a prediction (black) obtained by convolving proboscis extension epochs with a Ca^2+^ response function. Explained variance is indicated (*R*^2^). **(b)** Δ-F/F, normalized with respect to the neuron’s 90^th^%ile across the time-series, as a function of proboscis extension (PE) number within a PE train for ROIs 0 (solid boxes, filled circles) and 1 (dashed boxes, open circles). **(c)** Explained variance (*R*^2^) between Δ-F/F time-series and a prediction from convolving proboscis extension epochs with a Ca^2+^ response function and a time-window. Time-windows that maximize the correlation for ROIs 0 (solid line) and 1 (dashed line) are indicated (red circles). **(d)** Behavioral epochs are color-coded. Representative Δ-F/F time-series from two ROIs (green) are overlaid with a prediction (black) obtained by convolving proboscis extension epochs with a Ca^2+^ response function as well as a time window indicated in panel B (red circles). Explained variance is indicated (*R*^2^). **(e)** Standard deviation projection image for a SS31232-Gal4 nervous system expressing smFP and stained for GFP (green) and Nc82 (blue). Cell bodies are indicated (white asterisks). Scale bar is 40 µm. **(f)** Projection as in **e** but for one MCFO-expressing, traced neuron (black asterisks). The brain’s GNG (yellow) and VNC’s intermediate tectulum (green), and prothoracic leg neuromere (yellow) are color-coded. Scale bar is 40 µm. **(g, h)** Higher magnification projections of **(top)** brains and **(bottom)** VNCs for SS31232-Gal4 animals expressing **(g)** the stochastic label MCFO, or **(h)** the synaptic marker, syt:GFP (green), and tdTomato (red). Insets magnify dashed boxes. Indicated are cell bodies (asterisks), and bouton-like structures (white arrowheads). Scale bars for brain images are 10 µm. Scale bars for VNC images and insets are 20 µm and 2 µm, respectively.

Thus, PE-AN activity seemed to convey the temporal integration or counting of discrete events [41, 42]. Therefore, we next asked if PE-AN activity might be better predicted using a PE regressor that integrates the number of PEs within a given time window. Remarkably, by testing a variety of window sizes, we determined that the most accurate prediction of PE-AN dynamics could be obtained with an integration window of more than 10 s **(Figure 10C, red circles)**. This additional integration window made it possible to predict both the undershoot and overshoot of PE-AN activity at the start and end of PE trains, respectively (**Figure 10D**).

Temporal integration can be implemented using a line attractor model [43, 44] based on recurrently connected circuits. To explore the degree to which PE-AN might support an integration of PE events through recurrent interconnectivity, we examined PE-AN morphologies more closely. PE-AN cell bodies were located in the anterior T1 neuromere (**Figure 10E**). From there they projected dense neurites into the midline of the T1 neuromere (**Figure 10F**). Among these neurites, we observed puncta and syt:GFP expression consistent with presynaptic terminals **(Figure 10G,H, bottom)**. Their dense and highly overlapping arbors would be consistent with a mutual interconnectivity between PE-ANs. These putatively recurrent connections might enable the integration of PE events over tens-of-seconds. This integration might filter out sparse PE events associated with feeding and allow PE-ANs to only signal long PE trains that might be observed during deep rest-states [45]. These signals are conveyed to the brain’s GNG **(Figure 10G,H, top)**.

## 3 Discussion

Animals must be aware of their own behavioral states to accurately interpret sensory cues and select appropriate future actions. Here, we examined how this self-awareness might be conveyed to the brain by studying the activity and targeting of ascending neurons within the *Drosophila* motor system. Specifically, we addressed a number of fundamental questions (**Figure 1A**). First, to what extent do ANs encode the low-level movements of joint and legs, or high-level behavioral states like walking and grooming? Second, are individual AN encodings narrow (conveying one movement or behavior), or broad (conveying multiple movements or behaviors)? Third, to what extent do ANs target multiple or single brain regions? Fourth, do ANs that convey distinct signals also target distinct brain regions? Fifth, which characteristics of an AN’s patterning in the VNC are predictive of their encoding? Sixth, are ANs a simple feedforward relay of signals to the brain, or might they also contribute to computations within the VNC? To address these questions, we performed a large-scale functional and anatomical screen by leveraging a library of *Drosophila* sparsely expressing driver lines that target small sets of ANs as well as new experimental and computational tools for recording and quantifying neural activity in behaving animals.

### 3.1 Encoding of high-level behavioral states

We discovered that ANs functionally encode high-level behavioral states (**Figure 11A**), predominantly those related to self-motion like walking and resting. These encodings could be further distinguished as either broad (e.g., walk-ANs and foreleg-ANs), or narrow (e.g., turn-ANs and eye groom-ANs). Similarly, neurons in the vertebrate dorsal spinocerebellar tract have been shown to be more responsive to whole limb versus individual joint movements [46]. To compensate for the technical hurdle of relating relatively rapid joint movements to slow calcium indicator kinetics, we convolved joint angle time-series’ with a decay kernel that would maximize the explanatory power of our regression analyses. Additionally, we confirmed that potential issues related to the non-orthogonality of joint angles and leg movements would not obscure our ability to explain the variance of AN neural activity **(Figure S3)**. Our observation that eye groom-AN activity could be explained by movements of the forelegs gave us further confidence that, in principle, leg movement encoding could be detected (**Figure 2C**). Nevertheless, to further confirm the absence of low-level joint and leg movement encoding, future work could directly manipulate the joints and legs of restricted animals while recording AN activity [47]. Finally, we sometimes observed that the activity of putative walk-encoding ANs was not fully explained by our walking regressor, nor our turn analysis, (e.g., SS44270, overlaps with SS41605). This suggests that some ANs may encode other features of locomotion.

**Figure 11:**
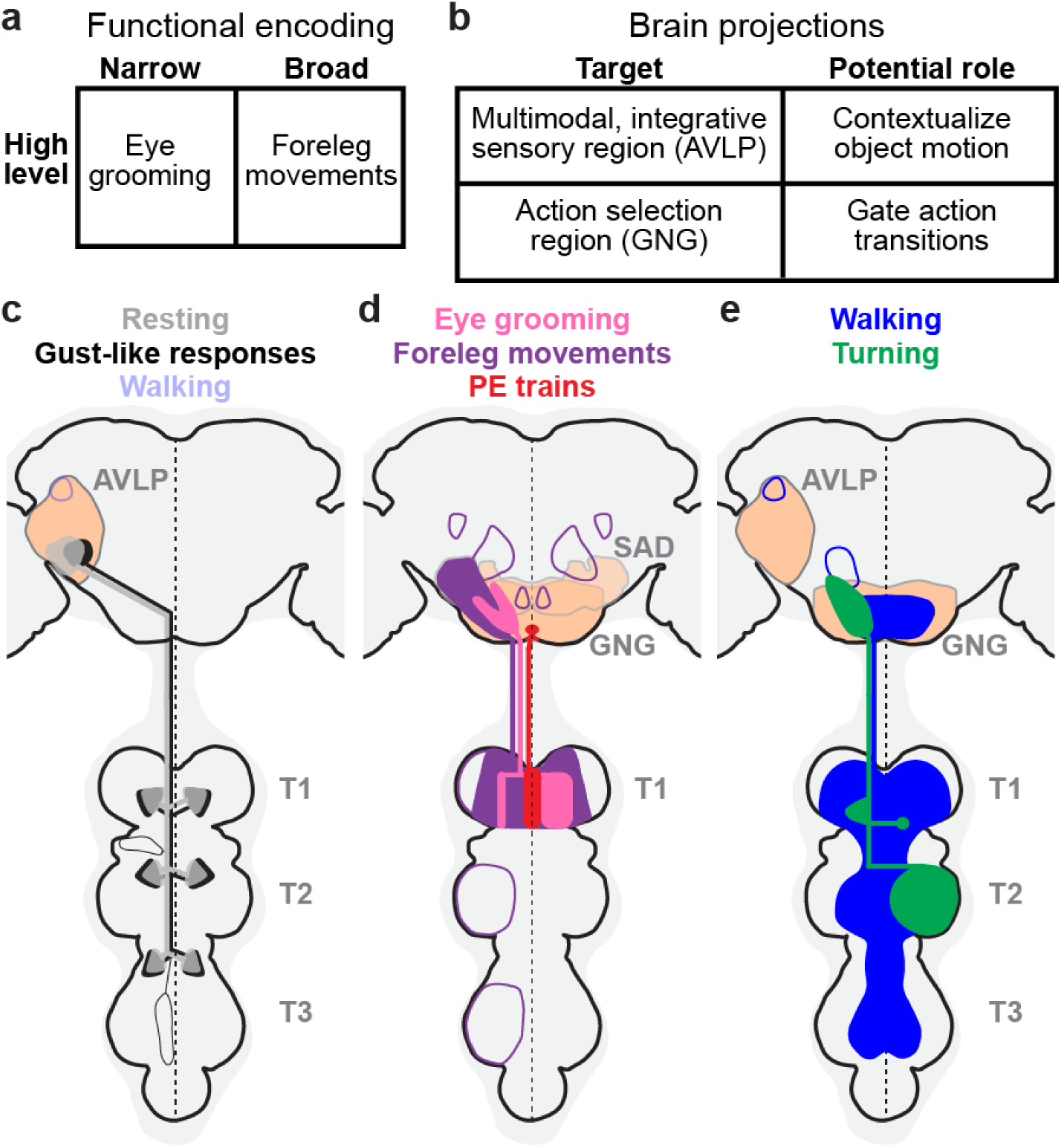
Summary of ascending neuron functional encoding, brain targeting, and VNC patterning. **(a)** Our functional screen shows that ANs encode high-level behaviors in a narrow (e.g., eye grooming), or broad (e.g., foreleg movements) manner. **(b)** Corresponding anatomical analysis shows that ANs primarily target the AVLP, a multimodal, integrative brain region, and the GNG, a region associated with action selection. **(c, d)** By comparing functional encoding with brain targeting and VNC patterning, we find that **(c)** signals critical for contextualizing object motion— walking, resting, and gust-like stimuli—are sent to the AVLP, while **(d)** signals indicating diverse ongoing behaviors are sent to the GNG, potentially to influence future action selection. **(e)** Broad (e.g., walking), or narrow (e.g., turning) behavioral encoding arises from diffuse and bilateral, or restricted and unilateral VNC innervations, respectively. **(c-e)** AN projections are color-coded by behavioral encoding. Axons and dendrites are not distinguished from one another. Brain and VNC regions are labelled. Frequently innervated brain regions—the GNG and AVLP—are highlighted (light orange). Less frequently innervated areas are outlined. The midline of the central nervous system is indicated (dashed line).

### 3.2 Predominant projection to the brain’s AVLP and GNG

We found that the vast majority of ANs do not project diffusely across the brain but rather specifically target either the AVLP and GNG (**Figure 11B**). We hypothesize that this may reflect the roles of behavioral state signals in two fundamental brain computations. First, the AVLP is a known site for multimodal, integrative sensory convergence [30–35]. Thus, the projection of ANs encoding resting, walking, and gust-like puffs to the AVLP (**Figure 11C**) may serve to contextualize time-varying visual and olfactory signals to indicate if they arise from self-motion, or from objects and odors moving in the world. A similar role of conveying self-motion has been proposed for neurons in the vertebrate dorsal spinocerebellar tract [16]. Second, the GNG is thought to be an action selection center [23, 36, 37]. Thus, the projection of ANs encoding diverse behaviors—walking, turning, foreleg movements, eye-grooming, and proboscis extensions (**Figure 11D,E**)—to the GNG may serve to indicate whether potential future actions are compatible with ongoing behaviors. This role would be consistent with hierarchical control approaches proposed in robotics [2]. Notably, walk-ANs that project to the ventral GNG may be neuromodulatory in nature. Thus, they may be well-poised to rapidly shift an animal’s internal state and the relative values of potential future actions.

Notably, the GNG is also heavily innervated by descending neurons (DNs). Because ANs and DNs both contribute to action selection [23,24,37,48], we speculate that they may connect within the GNG to form a feedback loop between the brain and motor system. Specifically, ANs that encode specific actions might excite DNs that drive the same actions, to generate behavioral persistence, and also suppress DNs that drive conflicting actions. For example, turn-ANs may excite DNa01 and DNa02 which control turning [27, 39, 49], and foreleg-ANs may excite aDN1 and aDN2 that control grooming [50]. Of course the opposite might also be true: ANs might *inhibit* DNs that encode the same action to ensure that motor actions are terminated once they have been performed. These competing hypotheses may soon be tested using emerging connectomics datasets [51].

### 3.3 Patterning within the VNC is predictive of behavioral encoding

The morphology of an AN’s neurites in the VNC are, to a remarkable degree, predictive of encoding (**Figure 11C-E**). We illustrate this in a few ways. First, ANs innervating all three leg neuromeres (T1, T2, and T3) encode global self-motion—walking, resting, and gust-like puffs. By contrast, those with more restricted projections to one neuromere (T1 or T2) encode discrete actions—turning, eye grooming, foreleg movements, and PEs. This might reflect the cost of neural wiring, a constraint that may encourage a neuron to sample the minimal sensory and motor information required to compute a particular behavioral state. Second, broadly tuned ANs (walking and foreleg-dependent behaviors) exhibited bilateral projections in the VNC while narrowly tuned ANs (turning and eye grooming) exhibited unilateral and smooth, putatively dendritic projections. This was correlated with the degree of synchrony in the activity of pairs of ANs **(Figure S8)**.

Strikingly, for all ANs that we examined in-depth, we found evidence of axon terminals within the VNC. Thus, ANs may not simply relay behavioral state signals to the brain but may also perform other roles. For example, they might contribute to motor control as components of central pattern generators (CPGs) that drive rhythmic movements [52], or rest-ANs might drive the muscle tone needed to maintain a natural resting posture. ANs might also participate in computing behavioral states. For example, we speculate that recurrent interconnectivity among PE-ANs might give rise to their temporal integration and encoding of PE number in a manner that can be modeled by line attractors [43, 44]. Finally, ANs might contribute to action selection within the VNC. For example, eye groom-ANs might project to the contralateral T1 neuromere to suppress circuits driving other foreleg-dependent behaviors like walking and foreleg rubbing.

### 3.4 Future work

Here we investigated animals that were generating spontaneous and puff-induced behaviors including walking and grooming. However, ANs likely also encode other behavioral states, unmeasured internal forces like posture-maintaining muscle stiffness, or even metabolic states. This is hinted at by the fact that the neural activity of some ANs were not well explained by any of our behavioral regressors (e.g., R87H02, R39G01, R69H10 and SS29633). Additionally, nearly one-third of the ANs we examined were unresponsive, possibly due to the lack of relevant context. In line with this, we found that some silent ANs could become very active during leg movements *only* when the spherical treadmill was removed (e.g., SS38631 and SS51017)**(Figure S9)**. Thus, future work should examine the encoding of ANs in a variety of contexts including tethered flight. Finally, it would also be interesting to test the degree to which AN encoding is genetically hardwired or capable of adapting following motor learning or after injury [53, 54]. In summary, here we have shown that ANs encode high-level behaviors that they convey to distinct integrative sensory and action selection centers in the brain. These findings can accelerate our understanding of how ascending neurons in the mammalian spinal cord influence decision-making in the brain [15, 16, 46, 55–57], and also inspire the development of more effective algorithms for robotic sensory contextualization and action selection [2].

## 5 Materials and Methods

### 5.1 Fly stocks

Split-Gal4 (spGal4) lines (SS*****) were generated by the Dickson laboratory and the FlyLight project (Janelia Research Campus, Ashburn VA USA; see **Table 1**). GMR lines, MCFO-5 (R57C10-Flp2::PEST in su(Hw)attP8;; HA-V5-FLAG), MCFO-7 (R57C10-Flp2::PEST in attP18;;HA-V5-FLAG-OLLAS) [26], and UAS-syt:GFP (Pw[+mC]=UAS-syt.eGFP1, w[*];;) were obtained from the Bloomington Stock Center. UAS-OpGCaM6f; UAS-tdTomato (; P20XUAS-IVS-Syn21-OpGCamp6F-p10 su(Hw)attp5; Pw[+mC]=UAS-tdTom.S3) was a gift from the Dickinson laboratory (Caltech, Pasadena CA USA). UAS-smFP (;; 10xUAS-IVS-myr::smGdP-FLAG (attP2)) was a gift from the McCabe laboratory (EPFL, Lausanne CH). Experimental animals were kept at 25*^◦^*C and 70% humidity on a 12-12 h day-light cycle.

**Table 2:** Activation (AD) and DNA-binding Domains (DBD) of split-Gal4 lines

|  | Driver line | AD | DBD |
| --- | --- | --- | --- |
| 1 | SS36131 | R70D06 | VT033054 |
| 2 | SS38592 | VT016458 | VT012410 |
| 3 | SS27485 | R75E01 | R18B05 |
| 4 | SS41822 | VT033054 | VT026646 |
| 5 | SS38624 | VT002081 | R85H01 |
| 6 | SS45605 | R15E01 | R41E03 |
| 7 | SS43652 | VT026477 | R38E07 |
| 8 | SS36112 | VT026646 | VT028606 |
| 9 | SS41806 | VT060737 | VT028606 |
| 10 | SS38631 | R72A10 | VT038208 |
| 11 | SS51029 | VT034810 | VT004985 |
| 12 | R85A11 | - | - |
| 13 | SS40489 | R36B06 | VT007767 |
| 14 | SS31480 | R68C10 | VT008150 |
| 15 | SS51021 | VT027767 | VT027005 |
| 16 | SS51017 | VT005404 | VT027767 |
| 17 | SS31456 | VT013500 | VT012768 |
| 18 | SS46233 | VT029814 | VT028464 |
| 19 | SS42749 | R66A06 | VT056770 |
| 20 | SS41815 | VT043377 | VT014669 |
| 21 | SS29633 | R33F06 | R76E11 |
| 22 | R87H02 | - | - |
| 23 | MAN | VT50660 | VT14014 |
| 24 | SS49172 | VT049120 | VT008188 |
| 25 | R36G04 | - | - |
| 26 | R39G01 | - | - |
| 27 | SS31219 | VT045153 | VT019074 |
| 28 | R30A08 | - | - |
| 29 | SS44270 | VT058560 | VT033054 |
| 30 | SS41605 | R80A11 | VT038205 |
| 31 | SS29579 | VT023828 | VT059224 |
| 32 | SS51046 | VT007177 | VT057280 |
| 33 | SS29893 | R67F03 | VT050658 |
| 34 | SS34574 | VT008537 | VT050658 |
| 35 | R70H06 | - | - |
| 36 | SS42740 | VT037865 | VT061717 |
| 37 | SS25469 | VT027704 | VT044958 |
| 38 | SS31232 | VT063643 | VT059781 |
| 39 | SS30303 | VT063643 | VT018278 |
| 40 | SS25451 | VT063643 | VT059224 |
| 41 | SS28596 | R94B04 | R86H08 |
| 42 | SS40134 | VT028320 | R49A01 |
| 43 | SS29621 | R22E07 | R30E10 |
| 44 | R69H10 | - | - |
| 45 | SS51038 | VT030558 | VT001497 |
| 46 | SS42008 | VT033469 | VT043682 |
| 47 | SS36118 | VT060737 | VT026477 |
| 48 | SS40619 | VT021853 | VT050234 |
| 49 | SS45363 | VT062587 | VT043920 |
| 50 | SS52147 | VT044164 | VT040034 |
| 51 | R38F09 | - | - |
| 52 | SS46269 | VT023490 | VT016254 |
| 53 | SS25470 | VT063643 | VT048352 |
| 54 | SS25478 | VT025966 | VT013121 |
| 55 | SS28382 | R18G02 | R49C03 |
| 56 | SS29574 | VT008660 | VT043400 |
| 57 | SS31899 | R26H04 | R46A10 |
| 58 | SS33380 | R19F01 | R60A06 |
| 59 | SS33433 | R94D12 | VT060731 |
| 60 | SS38012 | R48E02 | R93B07 |
| 61 | SS38386 | VT016966 | VT046334 |
| 62 | SS38687 | R30A02 | VT015159 |
| 63 | SS46290 | VT029750 | VT043288 |
| 64 | SS46300 | VT043146 | VT000254 |
| 65 | SS48406 | VT048352 | VT039769 |
| 66 | SS48409 | VT036302 | VT049125 |
| 67 | SS49982 | R77D08 | VT029514 |
| 68 | SS50004 | VT017645 | VT049348 |
| 69 | SS50013 | VT008992 | VT039485 |
| 70 | SS50652 | R60C01 | R80B01 |
| 71 | SS36132 | R70D06 | VT025996 |
| 72 | SS36133 | R70D06 | VT026646 |
| 73 | SS38598 | VT024634 | VT016458 |
| 74 | SS41808 | VT060737 | VT033054 |
| 75 | SS41809 | R20E05 | VT033054 |
| 76 | SS41820 | VT060737 | VT025996 |
| 77 | SS41821 | R20E05 | VT025996 |
| 78 | SS42007 | VT033469 | VT026646 |
| 79 | SS42707 | VT061717 | VT045101 |
| 80 | SS43528 | VT025966 | R93B07 |
| 81 | SS48632 | VT036302 | R93B07 |
| 82 | SS51024 | VT004985 | VT034810 |

**Table 2 continued from previous page**
|  | Driver line | AD | DBD |
| --- | --- | --- | --- |
| 83 | SS52108 | VT063231 | R69H06 |
| 84 | SS52106 | VT063231 | VT063626 |
| 85 | SS52107 | VT063231 | VT021731 |
| 86 | R86H08 | - | - |
| 87 | SS29889 | R64G04 | VT008537 |
| 88 | SS29890 | R64G04 | VT050658 |
| 89 | SS29605 | VT019902 | VT048942 |
| 90 | SS31246 | VT038171 | VT021780 |
| 91 | SS22721 | R92D09 | R92A07 |
| 92 | R75E01 | - | - |
| 93 | SS37652 | VT040698 | VT023490 |
| 94 | SS41602 | R75E01 | R74C01 |
| 95 | SS43651 | VT026477 | VT039361 |
| 96 | SS44305 | R21E09 | VT016966 |
| 97 | SS46255 | R24H02 | VT037862 |
| 98 | SS41824 | R20E05 | VT026646 |
| 99 | SS25488 | VT029593 | VT020527 |
| 100 | R81G07 | - | - |
| 101 | SS45635 | VT008882 | VT014208 |
| 102 | SS45648 | VT008808 | VT029814 |
| 103 | SS46290 | VT029750 | VT043288 |
| 104 | SS46847 | VT023490 | VT039485 |
| 105 | SS47868 | R24H02 | VT002064 |
| 106 | SS50282 | VT037554 | VT012768 |
| 107 | SS50829 | VT033290 | VT027767 |
| 108 | R88C08 | - | - |

### 5.2 *In vivo* two-photon calcium imaging experiments

Two-photon imaging was performed on 3-6 days post-eclosion (dpe) female flies as described in [27] with minor changes in the recording configuration. We imaged coronal sections of AN axons in the cervical connective to avoid having neurons move outside the field of view due to behavior-related tissue deformations. Imaging was performed using a Galvo-Galvo scanning system. Image dimensions ranged from 256 x 192 pixels (4.3 fps) to 320 x 320 pixels (3.7 fps), depending on the location of axonal regions-of-interest (ROIs) and the degree of displacement caused by animal behavior. During two-photon imaging, a 7-camera system was used to record fly behaviors as described in [28]. Rotations of the spherical treadmill, and the timing of puff stimuli were also recorded. Air or CO_2_ puffs (0.08 L/min) were controlled using either a custom Python script, or manually with an Arduino controller. Puffs were delivered through a syringe needle positioned in front of the animal to stimulate behavior in sedentary animals, or to interrupt ongoing behaviors. To synchronize signals acquired at different sampling rates—optic flow sensors, two-photon images, puff stimuli, and videography—signals were digitized using a BNC 2110 terminal block (National Instrument, USA) and saved using ThorSync software (Thorlabs, USA). Sampling pulses were then used as references to align data based on the onset of each pulse. Then signals were interpolated using custom Python scripts.

### 5.3 Immunofluorescence tissue staining and confocal imaging

Fly brains and VNCs from 3-6 dpe female flies were dissected and fixed as described in [27] with small modifications in staining including antibodies and incubation conditions (see details below). Both primary (rabbit anti-GFP at 1:500, Thermofisher RRID: AB 2536526; mouse anti-Bruchpilot / nc82 at 1:20, Developmental Studies Hybridoma Bank RRID: AB 2314866) and secondary antibodies (goat anti-rabbit secondary antibody conjugated with Alexa 488 at 1:500; Thermofisher, RRID: AB 143165; goat anti-mouse secondary antibody conjugated with Alexa 633 at 1:500; Thermofisher, RRID: AB 2535719) for smFP and nc82 staining were performed at room temperature for 24h.

To perform high-magnification imaging of MCFO samples, nervous tissues were incubated with primary antibodies: rabbit anti-HA-tag at 1:300 dilution (Cell Signaling Technology, RRID:AB 1549585), rat anti-FLAG-tag at 1:150 dilution (DYKDDDDK; Novus, RRID:AB 1625981), and mouse anti-Bruchpilot/nc82 at 1:20 dilution. These were diluted in 5% normal goat serum in PBS with 1% Triton-X (PBSTS) for 24 h at room temperature. The samples then were rinsed 2-3 times in PBS with 1% Triton-X (PBST) for 15 min before incubation with secondary antibodies: donkey anti-rabbit secondary antibody conjugated with AlexaFluor 594 at 1:500 dilution (Jackson ImmunoResearch Labs, RRID:AB 2340621), donkey anti-rat secondary antibody conjugated with AlexaFluor 647 at 1:200 dilution (Jackson ImmunoResearch Labs, RRID:AB 2340694), and donkey anti-mouse secondary antibody conjugated with AlexaFluor 488 at 1:500 dilution (Jackson ImmunoResearch Labs, RRID:AB 2341099). These were diluted in PBSTS for 24 h at room temperature. Again, samples were rinsed 2-3 times in PBS with 1% Triton-X (PBST) for 15 min before incubation with the last diluted antibody: rabbit anti-V5-tag (GKPIPNPLLGLDST) conjugated with DyLight 550 at 1:300 dilution (Cayman Chemical, 11261) for another 24 h at room temperature.

To analyze single neuron morphological patterning, we crossed spGal4 lines with MCFO-7 [26]. Dissection and MCFO staining were performed by Janelia FlyLight according to the FlyLight ‘IHC-MCFO’ protocol: https://www.janelia.org/project-team/flylight/protocols. Samples were imaged on an LSM710 confocal microscope (Zeiss) with a Plan-Apochromat 20x /0.8 M27 objective. To prepare samples expressing tdTomato and syt:GFP, we chose to only stain tdTomato to minimize false positive signals for the synaptotagmin marker. Samples were incubated with a diluted primary antibody: rabbit polyclonal anti-DsRed at 1:1000 dilution (Takara Biomedical Technology, RRID: AB 10013483) in PBSTS for 24 h at room temperature. After rinsing, samples were then incubated with a secondary antibody: donkey anti-rabbit secondary antibody conjugated with Cy3 (Jackson ImmunoResearch Labs, RRID:AB 2307443). Finally, all samples were rinsed 2–3 times for 10 min each in PBST after staining and then mounted onto glass slides with bridge coverslips in Slowfade mounting-media (Thermofisher, S36936).

Confocal imaging was performed as described in [27]. In addition, high-resolution images for visualizing fine structures were captured using a 40x oil-immersion objective lens with an NA of 1.3 (Plan-Apochromat 40x/1.3 DIC M27, Zeiss) on an LSM700 confocal microscope (Zeiss). The zoom factor was adjusted based on the ROI size of each sample between 84.23×84.23 *µ*m^2^ and 266.74×266.74 *µ*m^2^. For high-resolution imaging, z-steps were fixed at 0.33 *µ*m. Images were denoised, their contrasts were tuned, and standard deviation z-projections were generated using Fiji ( [58]).

### 5.4 Two-photon image analysis

Raw two-photon imaging data were converted to gray-scale *.tiff image stacks for both green and red channels using custom Python scripts. RGB image stacks were then generated by combining both image stacks in Fiji ( [58]). We used AxoID to perform ROI segmentation and to quantify fluorescence intensities. Briefly, AxoID was used to register images using cross-correlation and optic flow-based warping [27]. Then, raw and registered image stacks underwent ROI segmentation, allowing %Δ-F/F values to be computed across time from absolute ROI pixel values. Simultaneously, segmented RGB image stacks overlaid with ROI contours were generated. Each frame of these segmented image stacks was visually examined to confirm AxoID segmentation, or to perform manual corrections using the AxoID GUI. In these cases, manually corrected %Δ-F/F and segmented image stacks were updated.

### 5.5 Behavioral data analysis

To reduce computational and data storage requirements, we recorded behaviors at 30 fps. This is nearly the Nyquist frequency for rapid walking (up to 16 step cycles/s [59]).

3D joint positions were estimated using DeepFly3D [28]. Due to the amount of data collected, manual curation was not practical. We therefore classified points as outliers when the absolute value of any of their coordinates (x, y, z) was greater than 5 mm (much larger than the fly’s body size). Furthermore, we made the assumption that joint locations would only be incorrectly estimated for one of the three cameras used for triangulation. The consistency of the location across cameras could be evaluated using the reprojection error. To identify a camera with a bad prediction, we calculated the reprojection error only using two of the three cameras. The outlier was then replaced with the triangulation result of the pair of cameras with the smallest reprojection error. The output was further processed and converted to angles as described in [60].

We classified behaviors based on a combination of 3D joint angle dynamics and rotations of the spherical treadmill. First, to capture the temporal dynamics of joint angles, we calculated wavelet coefficients for each angle using 15 frequencies between 1 Hz and 15 Hz [61, 62]. We then trained a histogram gradient boosting classifier [63] using joint angles, wavelet coefficients, and ball rotations as features. Because flies perform behaviors in an unbalanced way (some behaviors are more frequenty than others), we balanced our annotations using SMOTE [64]. The model was validated using 5-fold, three times repeated, stratified cross-validation. Fly speeds and heading directions were estimated using optical flow sensors [27]. To further improve the accuracy of the onset of walking we applied empirically-determined thresholds (pitch: 0.0038; roll: 0.0038; yaw: 0.014) to the rotational velocities of the spherical treadmill. The rotational velocities were smoothed and denoised using a moving average filter (length 81). All frames that were not previously classified as grooming or pushing (and for which the spherical treadmill was classified as moving) were labeled as ‘walking’. These were furthered subdivided into forward or backward walking depending on the sign of the pitch velocity. Conversely, frames for which the spherical treadmill was not moving were labeled as ‘resting’. To reduce the effect of optical flow measurement jitter, walking and resting labels were processed using a hysteresis filter that only changes state if at least 15 consecutive frames are in a new state. Classification in this manner was generally effective but most challenging for kinematically similar behaviors like eye- and antennal-grooming, or hindleg rubbing and abdominal grooming **(Figure S10)**.

Proboscis extension (PE) events were classified based on the length of the proboscis **(Figure S2)**. First, we trained a deep network [38] to identify the tip of the proboscis and a static landmark (the ventral part of the eye) from side-view camera images. Then, the distance between the tip of proboscis and this static landmark was calculated to obtain the PE length for each frame. A semi-automated PE event classifier was made by first denoising the traces of PE distances using a median filter with a 0.3 s running average. Traces were then normalized to be between 0 (baseline values) and 1 (maximum values). Next, PE speed was calculated using a data point interval of 0.1 s to detect significant changes in PE length. This way, only peaks larger than a manually set threshold of 0.03 Δ-norm.length/0.1 s were considered. Because the peak speed usually occurred during the rising phase of a PE, a kink in PE speed was identified by multiplying the peak speed with an empirically-determined factor ranging from 0.4 to 0.6, and finding that speed within 0.5 s prior to the peak speed. The end of a PE was the time-point at which the same speed was observed within 2 s after the peak PE speed. This filtered out occasions where the proboscis remained extended for long periods of time. All quantified PE lengths and durations were then used to build a filter to remove false positives. PEs were then binarized to define PE epochs.

To quantify animal movements when the spherical treadmill was removed, we manually thresholded the variance of pixel values from a side view camera within a region of the image that included the fly. Pixel value changes were calculated using a running window of 0.2 s. Next, the standard deviation of pixel value changes was generated using a running window of 0.25 s. This trace was then smoothed and values lower than the empirically-determined threshold were called ‘resting’ epochs. The remainder were considered ‘movement’ periods.

### 5.6 Regression analysis of PE integration time

To investigate the integrative nature of the PE-AN responses, we convolved PE traces with uniform time windows of varying sizes. This convolution was performed such that the fluorescence at each time point would be the sum of fluorescence during the previous ‘window size’ frames (i.e., not a *centered* sliding window but one that only uses previous time points), effectively integrating over the number of previous PEs. This integrated signal was then masked such that all time points where the fly was not engaged in PE were set to zero. Then, this trace was convolved with a calcium indicator decay kernel, notably yielding non-zero values in the time intervals between PEs. We then determined the explained variance as described elsewhere and finally chose a window size maximizing the explained variance.

### 5.7 Linear modeling of neural fluorescence traces

Each regression matrix contains elements corresponding to the results of a ridge regression model for predicting the time-varying fluorescence 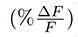 of ANs using specific regressors (e.g., low-level joint angles, or high-level behaviors). To account for slow calcium indicator decay dynamics, each regressor was convolved with a calcium response function. The half-life of the calcium response function was chosen from a range of 0.2 s to 0.95 s [65] in 0.05 s steps, in order to maximize the variance in fluorescence traces that convolved regressors could explain. The rise time was fixed at 0.1415 s [65]. The ridge penalty parameter was chosen using nested 10-fold stratified cross-validation [66]. The intercept and weights of all models were restricted to be positive, limiting our analysis to excitatory neural activity. Values shown in the matrices are the mean of 10-fold stratified cross-validation. We calculated Unique (UEV) and All-Explained Variance (AEV) by temporally shuffling the regressor in question, or all other regressors, respectively [4]. We tested the overall significance of our models using an F-statistic to reject the null hypothesis that the model does not perform better than an intercept-only model. The prediction of individual traces were performed using a single regressor plus intercept. Therefore they were not regularized.

### 5.8 Behavior-based neural activity analysis

For a given behavior, Δ-F/F traces were compiled, cropped, and aligned with respect to their onset times. Mean and 95% confidence intervals for each time point were then calculated from these data. Because the duration of each behavioral epoch was different, we only computed mean and confidence intervals for epochs that had at least 5 data points.

To test if each behavior-triggered average Δ-F/F was significantly different from the baseline, first, we aligned and upsampled fluorescence data that were normalized between 0 (baseline mean) and 1 (maximum) for each trial. For each behavioral epoch, the first 0.7 s of data were removed. This avoided contaminating signals with neural activity from preceding behaviors (due to the slow decay dynamics of OpGCaMP6f). Next, to be conservative in judging whether data reflected noisy baseline or real signals, we studied their distributions. Specifically, we tested the normality of twenty resampled groups of 150 bootstrapped datapoints—a size that reportedly maximizes the power of the Shapiro-Wilk test [67]. If a majority of results did not reject the null hypothesis, the entire recording was considered baseline noise and the Δ-F/F for a given behavioral class was not considered significantly different from baseline. On the other hand, if the datapoints were not normally distributed, the baseline was determined using an Otsu filter. For recordings that passed this test of normality, if the majority of six ANOVA tests on the bootstrapped data rejected the null hypothesis and the datapoints of a given behavior were significantly different (\*\*\**p<*0.001, \*\**p<*0.01, \**p<*0.05) from baseline (as indicated by a posthoc Tukey test), these data were considered signal and not noise.

To analyze PE-AN responses to each PE during PE trains, putative trains of PEs were manually identified to exclude discrete PE events. PE trains included at least 3 consecutive PEs in which each PE lasted at least 1 s and there was less than 3 s between each PE. Then, the mean fluorescence of each PE was computed for 25 PE trains (n=11 animals). The median, IQR, and 1.5 IQR were then computed for PEs depending on their ordered position within their PE trains. We focused our analysis on the first 11 PEs because they had a sufficiently large amount of data.

### 5.9 Neural fluorescence-triggered averaging of spherical treadmill rotational velocities

A semi-automated neural fluorescence event classifier was constructed by first denoising Δ-F/F traces by averaging them using a 0.6 s running window. Traces were then normalized to be between 0 (their baseline values) and 1 (their maximum values). To detect large deviations, the derivative of the normalized Δ-F/F time-series was calculated at an interval of 0.1 s. Only peaks greater than an empirically determined threshold of 0.03 *d* norm Δ-F/F / 0.1 s were considered events. Because peak fluorescence derivatives occurred during the rising phase of neural fluorescence events, the onset of a fluorescence event was identified as the time where the Δ-F/F derivative was 0.4-0.6x the peak within the preceding 0.5 s time window. The end of the event was defined as the time that the Δ-F/F signal returned to the amplitude at event onset before the next event. False positives were removed by filtering out events with amplitudes and durations that were lower than the empirically determined threshold. Neural activity event analysis for turn-ANs was performed by testing if the mean normalized fluorescence event for one ROI was larger than the other ROI by an empirically determined factor of 0.2x. Corresponding ball rotations for events that pass this criteria were then collected. Fluorescence events onsets were then set to 0 s and aligned with spherical treadmill rotations. Using these rotational velocity data, we calculated the mean and 95% confidence intervals for each time point with at least five data points. A 1 s period before each fluorescence event was also analyzed as a baseline for comparison.

### 5.10 Brain and VNC confocal image registration

All confocal images, except for MCFO image stacks, were registered based on nc82 neuropil staining. We built a template and registered images using the CMTK munger extension [68]. Code for this registration process can be found at: https://github.com/NeLy-EPFL/MakeAverageBrain/tree/workstation. Brain and VNC images were registered to JRC 2018 templates [69] using the Computational Morphometry Toolkit: https://www.nitrc.org/projects/cmtk. The template brain and VNC can be downloaded here: https://www.janelia.org/open-science/jrc-2018-brain-templates.

### 5.11 Analysis of individual AN innervation patterns

Single AN morphologies were traced by masking MCFO confocal images using either active tracing, or manual background removal in Fiji [58]. Axons in the brain were manually traced using the Fiji plugin ‘SNT’. Most neurites in the VNC were isolated by (i) thresholding to remove background noise and outliers, and (ii) manually masking debris in images. In the case of ANs from SS29579, a band-pass color filter was applied to isolate an ROI that spanned across two color channels. The boundary of the color filter was manually tuned to acquire the stack for a single neuron mask. After segmentation, the masks of individual neurons were applied across frames to calculate the intersectional pixel-wise sum with another mask containing either (i) neuropil regions of the brain and VNC, (ii) VNC segments, or (iii) left and right halves of the VNC. Brain and VNC neuropil regions and their corresponding abbreviations were according to established nomenclature [70]. Neuropil region masks can be downloaded here: https://v2.virtualflybrain.org. These were also registered to the JRC 2018 template. Masks for T1, T2, and T3 VNC segments were based on previously delimited boundaries [37]. The laterality of a neuron’s VNC innervation was calculated as the ratio of the absolute difference between its left and right VNC innervations divided by its total innervation. Masks for the left and right VNC were generated by dividing the VNC mask across its midline.

### 5.12 AxoID: a deep learning-based software for tracking axons in imaging data

AxoID aims to extract the GCaMP fluorescence values for axons present on coronal section two-photon microscopy imaging data. In this manuscript, it is used to record activity from ascending neurons (ANs) passing through the *Drosophila melanogaster* cervical connective. Fluorescence extraction works by performing the following three main steps **(Figure S11A)**. First, during a *detection* stage, ROIs corresponding to axons are segmented from images. Second, during a *tracking* stage, these ROIs are tracked across frames. Third, *fluorescence* is computed for each axon over time.

To track axons, we used a two-stage approach: detection and then tracking. This allowed us to improve each problem separately without the added complexity of developing a detector that must also do tracking. Additionally, this allowed us to detect axons without having to know how many there are in advance. Lastly, significant movement artifacts between consecutive frames pose additional challenges for robustness in temporal approaches while, in our case, we can apply the detection on a frame-by-frame basis. However, we note that we do not leverage temporal information.

#### 5.12.1 Detection

Axon detection consists of finding potential axons by segmenting the background and foreground of each image. An ROI or putative axon is defined as a group of connected pixels segmented as foreground. Pixels are considered connected if they are next to one another.

By posing detection as a segmentation problem, we have the advantage of using standard computer vision methods like thresholding, or artificial neural networks that have been developed for medical image segmentation. Nevertheless, this simplicity has a drawback: if axons appear very close to one another and their pixels are connected, they may be segmented as one ROI rather than two. We try to address this issue using an ROI separation approach described later.

Image segmentation is performed using deep learning on a frame-by-frame basis, whereby a network generates a binary segmentation of a single image. As a post-processing step, all ROIs smaller than a minimum size are discarded. Here, we empirically chose 11 pixels as the minimum size as a trade-off between removing small spurious regions while still detecting small axons.

We chose to use a U-Net model [71] with slight modifications because of its, or its derivatives’, performance on recent biomedical image segmentation problems [72–74]. We add zero-padding to the convolutions to ensure that the output segmentation has the same size as the input image, thus fully segmenting it in a single pass, and modify the last convolution to output a single channel rather than two. Batch normalization [75] is used after each convolution and its non-linearity function. Finally, we reduce the width of the network by a factor of 4: each feature map has 4 times fewer channels than the original U-Net, not counting the input or output. The input pixel values are normalized to the range [-1, 1], and the images are sufficiently zero-padded to ensure that the size can be correctly reduced by half at each max-pooling layer.

To train the deep learning network, we use the Adam optimizer [76] on the binary cross-entropy loss with weighting. Each background pixel is weighted based on its distance to the closest ROI, given by 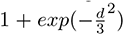 with *d* the Euclidean distance, plus a term that increases if the pixel is a border between two axons, given by 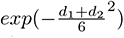 with *d*_1_ and *d*_2_ as the distances to the two closest ROIs, as in [71]. These weights aim to encourage the network to correctly segment the border of the ROI and to keep a clear separation between two neighboring regions. At training time, the background and foreground weights are scaled by 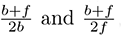, respectively, to take into account the imbalance in the number of pixels, where *b* and *f* are the quantity of background and foreground (i.e., ROI) pixels in the entire training dataset. To evaluate the resulting deep network, we use the Sørensen-Dice coefficient [77, 78] at the pixel level, which is equivalent to the F1-score. The training is stopped when the validation performance does not increase anymore.

The network was trained on a mix of experimental and synthetic data. We also apply random gamma corrections to the training input images, with *γ* sampled in [0.7, 1.3] to keep reasonable values, and to encourage robustness against intensity variations between experiments. The target segmentation of the axons on the experimental data was generated with conventional computer vision methods. First, the images were denoised with the non-local means algorithm [79] using the Python implementation of OpenCV [80]. We used a temporal window size of 5, and performed the denoising separately for the red and green channels, with a filter strength *h* = 11. The grayscale result was then taken as the per-pixel maximum over the channels. Following this, the images were smoothed with a Gaussian kernel of standard deviation 2 pixels, and thresholded using Otsu’s method [81]. A final erosion was applied and small regions below 11 pixels were removed. All parameter values were set empirically to generate good qualitative results. In the end, the results were manually filtered to keep only data with satisfactory segmentation.

Because the experimental data have a fairly simply visual structure, we constructed a pipeline in Python to generate synthetic images visually similar to real ones. This was achieved by first sampling an image size for a given synthetic experiment, then by sampling 2D Gaussians over it to simulate the position and shape of axon cross-sections. After this, synthetic tdTomato levels were uniformly sampled and GCaMP dynamics were created for each axons by convolving a GCaMP response kernel with Poisson noise to simulate spikes. Then, the image with the Gaussian axons was deformed multiple times to make different frames with artificial movement artifacts. Eventually, we sampled from the 2D Gaussians to make the axons appear pixelated, and added synthetic noise to the images. In the end, we chose a deep learning-based approach because our computer vision pipeline alone was not be robust enough. Our pipeline is used to generate a target segmentation dataset from which we manually select a subset of acceptable results. These results are then used to train the deep learning model.

##### Fine-tuning

At the beginning of the detection stage, an optional fine-tuning of the network can be applied to try to improve the segmentation of axons. The goal is to have a temporary network adapted to the current data for better performance. To do this, we train the network on a subset of experimental frames using automatically generated target segmentations.

The subset of images is selected by finding a cluster of frames with high cross-correlation-based similarity. For this, we only consider the tdTomato channel to avoid the effects of GCaMP dynamics. Each image is first normalized by its own mean pixel intensity *µ* and standard deviation *σ*: *p*(*i, j*) *←* 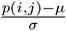, where *p*(*i, j*) is the pixel intensity *p* at the pixel location *i, j*. The cross-correlation is then computed between each pair of normalized images *p_m_* and *p_n_* as 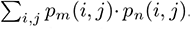. Afterwards, we take the opposite of the cross-correlation as a distance measure and use it to cluster the frames with the OPTICS algorithm [82]. We set the minimal number of sample for a cluster to 20, in order to maintain at least 20 frames for fine-tuning, and a maximum neighborhood distance of half the largest distance between frames. Finally, we select the cluster of images with the highest average cross-correlation (i.e., the smallest average distance between its elements).

Then, to generate a target segmentation image for these frames, we take their temporal average and optionally smooth it, if there are less than 50 images, to help remove the noise. The smoothing is done by filtering with a Gaussian kernel of standard deviation 1 pixel, then median filtering over each channel separately. The result is then thresholded through a local adaptive method, computed by taking the weighted mean of the local neighborhood of the pixel, subtracted by an offset. We apply Gaussian weighting over windows of 25 *×* 25 pixels, with an offset of *−*0.05, determined empirically. Finally, we remove regions smaller than 11 pixels. The result serves as a target segmentation image for all of the fine-tuning images.

The model is then trained on 60% of these frames with some data augmentation, while the other 40% are used for validation. The fine-tuning stops automatically if the performance on the validation frames drops. This avoids bad generalization for the rest of the images. The binary cross-entropy loss is used, with weights computed as discussed previously. For the data augmentation, we use random translation (±20%), rotation (±10°), scaling (±10%), and shearing (±5°).

#### 5.12.2 Tracking

Once the regions of interest are segmented, the next step of the pipeline consists of tracking the axons through time. This means defining which axons exist, and then finding the ROI they correspond to in each frame.

##### Tracking template

To accomplish this, the tracker records the number of axons, their locations with respect to one another, and their areas. It stores this information into what we call the ‘tracker template’. Then, for each frame, the tracker matches its template axons to the ROIs to determine which regions correspond to which axons.

The tracker template is built iteratively. It is first initialized and then updated by matching with all experimental data. The initialization depends on the optional fine-tuning in the detection step. If there is fine-tuning, then the smoothed average of the similar frames and its generated segmentation are used. Otherwise, one frame of the experiment is automatically selected. For this, AxoID considers only the frames with a number of ROIs equal to the most frequent number of ROIs, and then selects the image with the highest cross-correlation with the temporal average of these frames. It is then smoothed and taken with the segmentation produced by the detection network as initialization. The cross-correlation and smoothing are computed identically as in the fine-tuning. Each ROI in the initialization segmentation defines an axon in the tracker template, with its area and position recorded as initial properties.

Afterwards, we update them by matching each experimental frame to the tracker template. It consists of assigning the ROI to the tracker axons, and then using these regions’ areas and positions to update the tracker. The images are matched sequentially, and the axons properties are taken as running averages of their matched regions. For example, considering the *n^th^* match, the area of an axon is updated as:

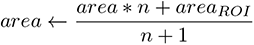

Because of this, the last frames are matched to a tracker template that is different from the one used for the first frames. Therefore, we fix the axons properties after the updates and match each frame again to obtain the final identities of the ROIs.

##### Matching

To assign axon identities to the ROIs of a frame, we perform a matching between them as discussed above. To solve it, we define a cost function for matching a template axon to a region which represents how dissimilar they are. Then, using the Hungarian assignment algorithm [83], we find the optimal matching with the minimum total cost **(Figure S11B)**.

Because some ROIs in the frame may be wrong detections, or some axons may not be correctly detected, the matching has to allow for the regions and axons to end up unmatched for some frames. Practically, we implement this by adding “dummy” axons to the matching problem with a flat cost. To guarantee at least one real match, the flat cost is set to the maximum between a fixed value and the minimum of the costs between regions and template axons with a margin of 10%: *dummy* = *max*(*v,* 1.1*· min*(costs)) with *v* = 0.3 the fixed value. Then, we can use the Hungarian method to solve the assignment, and all ROIs linked to these dummy axons can be considered unmatched.

We define the cost of assigning a frame’s ROI *i* to a tracker template axon *k* by their absolute difference in area plus the mean cost of an optimal inner matching of the other ROI to the other axons assuming *i* and *k* are already matched:

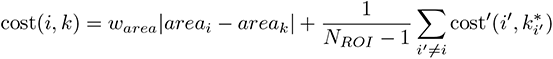

where *w_area_* = 0.1 is a weight for balancing the importance of the area, *N_ROI_* is the number of ROI in the frame, and 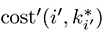 is the inner cost of assigning region 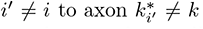 selected in an “inner” assignment problem, see below. In other words, the cost is relative to how well the rest of the regions and axons match if we assume that *i* and *k* are already matched.

The optimal inner matching is computed through another Hungarian assignment, for which we define another cost function. For this “inner” assignment problem, the cost of matching an ROI 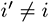 and a template axon 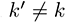 is defined by how far they are and their radial difference with respect to the matched *i* and *k*, plus their difference in area:

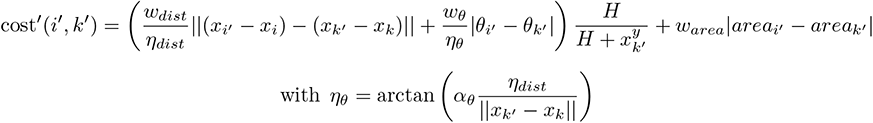

where *w_dist_* = 1.0, *w_θ_* = 0.1, and *w_area_* = 0.1 are weighting parameters, *η_dist_* = min(*H, W*) and *η_θ_* are normalization factors with *H* and *W* the height and width of the frame and *α_θ_* = 0.1 a secondary normalization factor. The *· ^y^* operation returns the height component of a vector, and the 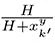 term is useful to reduce the importance of the first terms if the axon *k^’^* is far from axon *k* in the height direction. This is needed as the scanning of the animal’s cervical connective is done from top to bottom, thus we need to allow for some movement artifacts between the top and bottom of the image. Note that the dummy axons for unmatched regions are also added to this inner problem.

This inner assignment is solved for each possible pair of axon-ROI to get all final costs. The overall matching is then performed with them. Because we are embedding assignments, the computational cost of the tracker increases exponentially with the number of ROIs and axons. It stays tractable in our case as we generally deal with few axons at a time. All parameter values used in the matching were found empirically by trial and error.

##### Identities post-processing: ROI separation

In the case of fine-tuning at the detection stage, AxoID will also automatically try to divide ROIs that are potentially two or more separate axons. We implement this to address the limitation introduced by detecting axons as a segmentation: close or touching axons may get segmented together.

To do this, it first searches for potential ROIs to be separated by reusing the temporal average of the similar frames used for the fine-tuning. This image is initially segmented as described before. Then local intensity maxima are detected on a grayscale version of this image. To avoid small maxima due to noise, we only keep those with an intensity *≥* 0.05, assuming normalized grayscale values in [0, 1]. Following this, we use the watershed algorithm, with the scikit-image [84] implementation, to segment the ROI based on the gray level and detected maxima. In the previous stages, we discarded ROIs under 11 pixels to avoid small spurious detections. Similarly, here we fuse together adjacent regions that are under 11 pixels to only output results after the watershedding above or equal to that size. Finally, a border of 1 pixel width is inserted between regions created from the separation of an ROI.

These borders are the divisions separating the ROI, referred to as “cuts”. We parameterize each of these as a line, defined as its normal vector **n** and distance *d* to the origin of the image (top-left). To report them on each frame, we first normalize this line to the current ROI, and then reverse that process with respect to the corresponding regions on the other frames. To normalize the line to an ROI, we fit an ellipse on the ROI contour in a least-square sense. Then the line parameters are transformed into this ellipse’s local coordinates following Algorithm 1. It is essentially like transforming the ellipse into a unit circle, centered and axis-aligned, and applying a similar transformation to the cutting line **(Figure S11C, middle)**. The choice of fitting an ellipse is motivated by the visual aspect of the axons in the experimental data as they are fairly similar to elongated ellipses. Considering this, a separation between two close ellipses could be simplified to a linear border, motivating the linear representation of the ROI separation.

**Algorithm 1:**
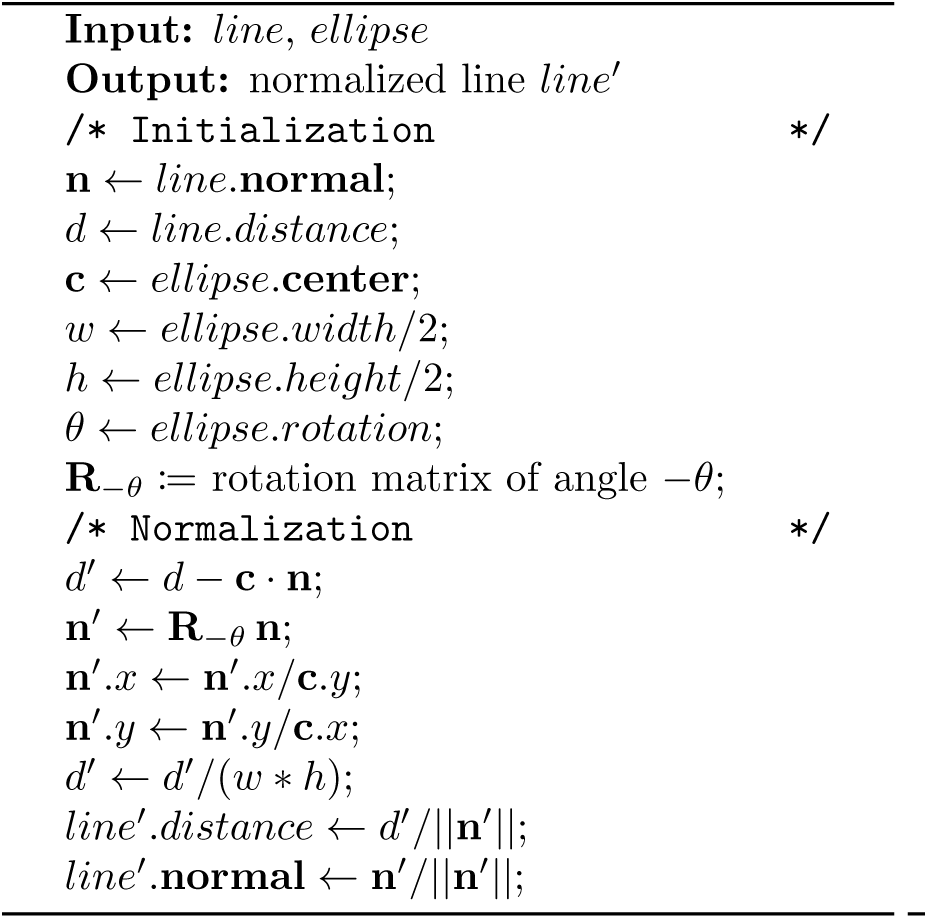
Normalize a line with an ellipse

**Algorithm 2:**
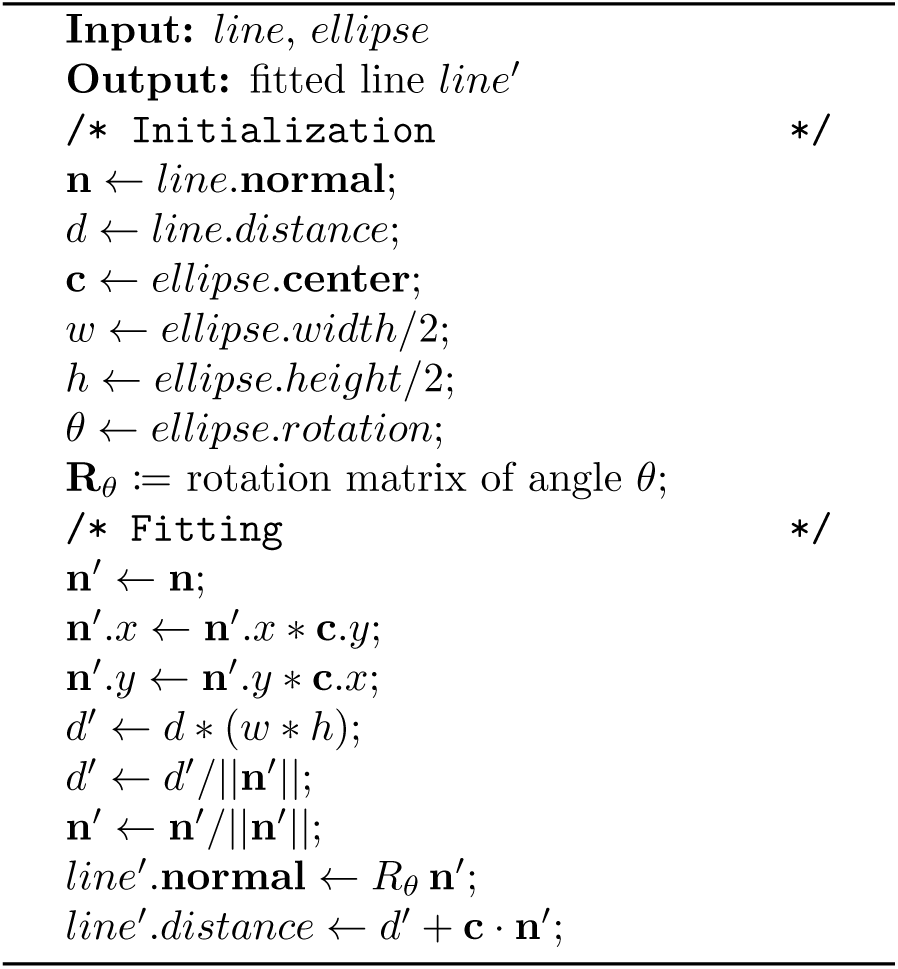
Fit a line to an ellipse

Because this is done as a post-processing step following tracking, we can apply that division on all frames. To do this, we again fit an ellipse to their ROI contours in the least-squares sense. Then, we take the normalized cutting line and fit it back to each of them according to Algorithm 2. This is similar to transforming the normalized unit circle to the region ellipse and applying the same transform to the line **(Figure S11C, right)**.

Finally, a new axon is defined for each cut. In each frame, the pixels of the divided region on the furthest side of the linear separation (with respect to the fitting ellipse center) are taken as the new ROI of that axon for that given frame.

In case there are multiple cuts of the same ROI (e.g., because three axons were close), the linear separations are ordered by distance to the center of the fitting ellipse and are then applied in succession. This is simple and efficient, but assumes there is little to no crossing between linear cuts.

#### 5.12.3 Fluorescence extraction

With the detection and tracking results, we know where each axon is in the experimental data. Therefore, to compute tdTomato and GCaMP fluorophore time-series we take the average of non-zero pixel intensities of the corresponding regions in each frame. We report the GCaMP fluorescence at time *t* as *F_t_*, and the ratio of GCaMP to tdTomato fluorescence at time *t* as *R_t_* to gain robustness against image intensity variations.

The final GCaMP fluorescence is reported as in [27]:

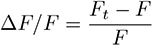

where *F* is a baseline fluorescence. Similarly, we report the ratio of GCaMP over tdTomato as in [27, 85]:

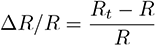

where *R* is the baseline. The baseline fluorescences *F* and *R* are computed as the minimal temporal average over windows of 10 s of the fluorophore time series *F_t_* and *R_t_*, respectively. Note that axons can be missing in some frames. For instance, if they were not detected or leave the image during movement artifacts. In this case, the fluorescence of that axon will have missing values at the time index *t* in which it was absent.

### 5.13 Overall workflow

To improve the performance of AxoID, the fluorescence extraction pipeline is applied three times: once over the raw data, once over the data registered using cross-correlation, and once over the data registered using optic-flow warping. Note that the fine-tuning in the detection stage is not used with the raw experimental data as it is based on the cross-correlation between the frames and would therefore lead to worse or redundant results with the data registered using cross-correlation. Eventually, the three fluorescence results can be visualized, chosen from, and corrected by a user through a GUI **(Figure S11D)**.

#### 5.13.1 Data registration

Registration of the experimental frames consists in transforming each image to make them similar to a reference image. The goal is to reduce the artifacts introduced by animal movements and to align axons across frames. This should help to improve the results of the detection and tracking.

##### Cross-correlation

Cross-correlation registration consists of translating an image so that its correlation to a reference is maximized. Note that the translated image wraps-around (e.g., pixels disappearing to the left reappear on the right). This aims to align frames against translations, but is unable to counter rotations or local deformations. We used the single step Discrete Fourier Transform (DFT) algorithm [86] to find the optimal translation of the frame. It first transforms the images into the Fourier domain, computes an initial estimate of the optimal translation, and then refines this result using a DFT. We based our Python implementation on previous work [87].

For each experiment, the second frame is taken as the reference frame to avoid recording artifacts that sometimes appear on the first recorded image.

##### Optic-flow registration

Optic flow-based registration was previously published [27]. Briefly, this approach computes an optic flow from the frame to a reference image, then deforms it by moving the pixels along that flow. The reference image is taken as the first frame of the experiment. This method has the advantage of being able to compute local deformations, but at a high computational cost.

#### 5.13.2 AxoID GUI

Finally, AxoID contains a GUI where a user can visualize the results, select the best one, and manually correct it.

First, the user is presented with three outputs of the fluorescence extraction pipeline from the raw and registered data with the option of visualizing different information to select the one to keep and correct. Here, the detection and tracking outputs are shown, as well as other information like the fluorescence traces in Δ*F/F* or Δ*R/R*. One of the results is then selected and used throughout the rest of the pipeline.

Following this, the user can edit the tracker template, which will then automatically update the ROI identities across frames. The template and the identities for each frame are shown, with additional information like the image used to initialized the template. The user has access to different tools: axons can be fused, for example, if they actually correspond to a single real axon that was incorrectly detected as two, and, conversely, one axon can be manually separated in two if two close ones are detected together. Moreover, useless axons or wrong detections can be discarded.

Once the user is satisfied with the overall tracker, they can correct individual frames. At this stage, it is possible to edit the detection results by discarding, modifying, or adding ROIs onto the selected image. Then, the user may change the tracking results by manually correcting the identities of these ROIs. In the end, the final fluorescence traces are computed on the selected outputs including user corrections.

## 8 Supplementary Videos

**Video 1: High-level behaviors, their associated 3D poses, and spherical treadmill rotational velocities.** Behaviors were captured from six camera views. Illuminated text (top) indicates the regressor being illustrated. Also shown are corresponding 3D poses (bottom-left) and spherical treadmill rotational velocities, proboscis extension (PE) lengths, and puff stimulation periods (bottom-right). https://www.dropbox.com/s/xed6jfgyqf7ubft/Video1.mov?dl=0

**Video 2: Representative data for 50 comprehensively analyzed, AN-targeting sparse driver lines.** Shown are: (**a**) spFP staining, (**b**) a representative two-photon microscope image, (**c**) outline of the associated cervical connective after filling the surrounding bath with fluorescent dye, (**d**) and PE length, puff stimuli, spherical treadmill rotational velocities, and AN (ROI) *6*F/F traces. Indicated above are regressors for forward walking (‘F.W.’), backward walking (‘B.W.’), resting (‘Rest’), eye grooming (‘Eye groom’), antennal grooming (‘Ant. groom’), foreleg rubbing (‘Fl. rub’), abdominal grooming (‘Abd. groom’), hindleg rubbing (‘Hl. rub’), and proboscis extension (‘PE’). For each driver line, the title indicates ‘date-Gal4-reporters-fly-trial’. https://www.dropbox.com/s/73aymyw3quiw142/Video2.mov?dl=0

**Videos 3 - 52: Representative behavioral videos and AN two-photon imaging data for 50 comprehensively analyzed, AN-targeting sparse driver lines.** https://drive.switch.ch/index.php/s/Q9K5BvugJc190rV

## 9 Data and code availability

Data are available at: https://dataverse.harvard.edu/dataverse/AN

Analysis code is available at: https://github.com/NeLy-EPFL/Ascending_neuron_screen_analysis_pipeline

AxoID code is available at: https://github.com/NeLy-EPFL/AxoID

## 10 Funding

PR acknowledges support from an SNSF Project grant (175667) and an SNSF Eccellenza grant (181239). FA acknowledges support from a Boehringer Ingelheim Fonds PhD stipend.

## 11 Acknowledgments

We thank the Janelia Research Campus FlyLight project for generating Ascending Neuron split-Gal4 driver lines.

## 12 Author Contributions

C-L.C. - Conceptualization, Methodology, Software, Validation, Formal Analysis, Investigation, Data Curation, Validation, Writing – Original Draft Preparation, Writing – Review & Editing, Visualization.

F.A - Methodology, Software, Formal Analysis, Investigation, Data Curation, Validation, Data Curation, Writing – Original Draft Preparation, Writing - Review & Editing.

R.M. - Methodology, Investigation, Data Curation, Validation. Writing - Review & Editing

V.M. - Investigation, Data Curation, Visualization. Writing - Review & Editing

N.T. - Methodology, Software, Formal Analysis, Data Curation Visualization. Writing - Review &

Editing

S.G. - Methodology, Software, Formal Analysis, Data Curation, Visualization. Writing - Review & Editing

B.D. - Resources, Writing - Review & Editing, Supervision, Project Administration, Funding Acquisition. Writing - Review & Editing

P.R. - Conceptualization, Methodology, Resources, Writing – Original Draft Preparation, Writing - Review & Editing, Supervision, Project Administration, Funding Acquisition.

## 13 Competing interests

The authors declare that no competing interests exist.

---

**Figure S1:**
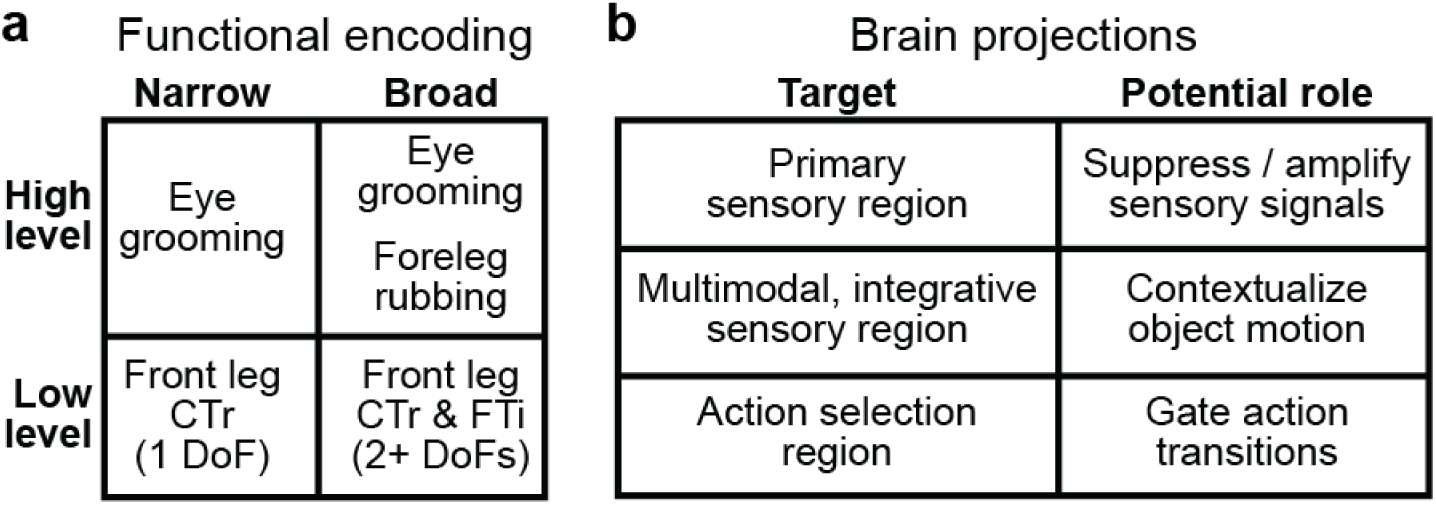
Hypothetical ascending neuron functional encoding and brain targeting. (**a**) ANs might encode high-level behaviors, or low-level limb kinematics. This encoding may be either narrow (e.g., one behavior, or joint degree-of-freedom), or broad (e.g., several behaviors, or joint DoFs). (**b**) ANs might target the brain’s (i) primary sensory regions (e.g., optic lobe, or antennal lobe) to perform sensory gain control, (ii) multimodal and integrative sensory regions (e.g., anterior ventrolateral protocerebrum, or mushroom body) to contextualize time-varying sensory cues, or (iii) action selection centers (e.g., gnathal ganglion, or central complex) to gate action transitions. Individual ANs may project broadly to multiple brain regions, or narrowly to one region.

**Figure S2:**
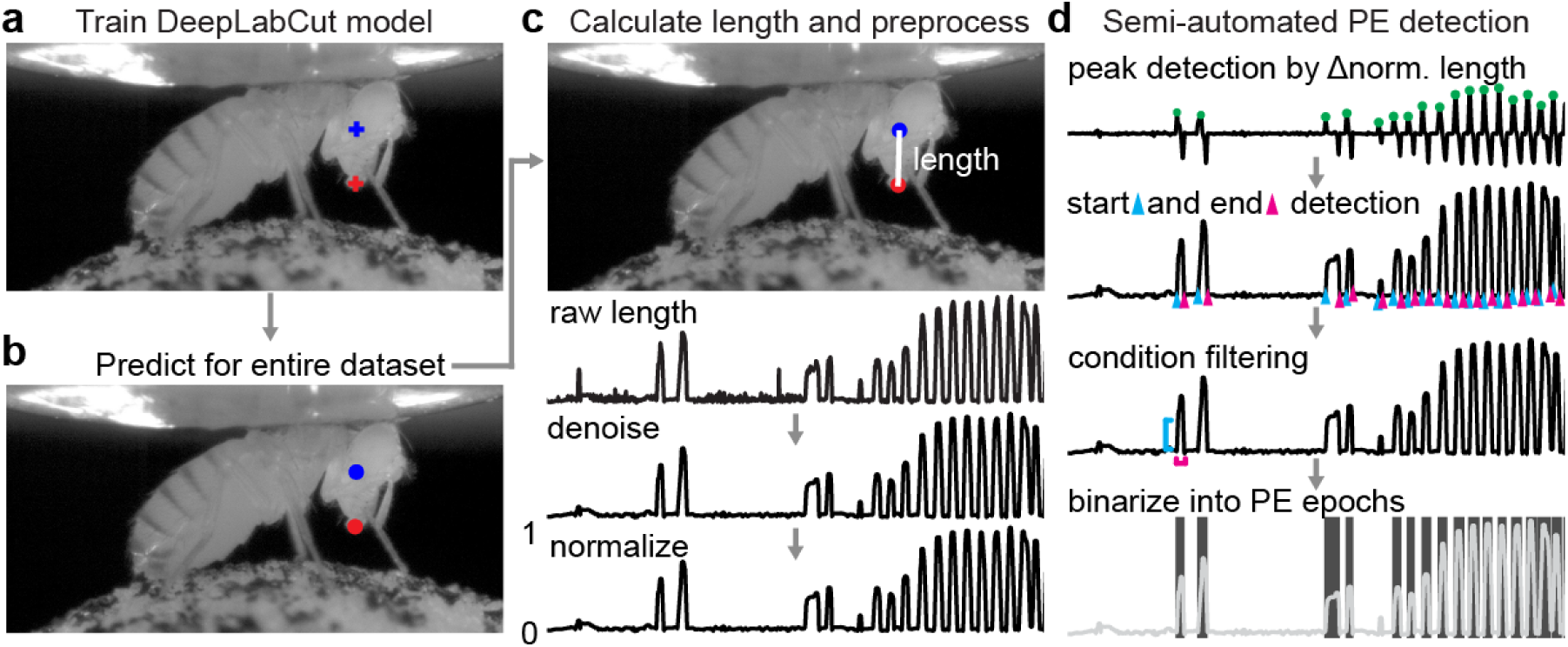
Semi-automated tracking of proboscis extensions. We detected proboscis extensions using side-view camera images. (**a**) First, we trained a deep neural network model with manual annotations of landmarks on the ventral eye (blue cross) and distal proboscis tip (red cross). (**b**) Then we applied the trained model to estimate these locations throughout the entire dataset. (**c**) Proboscis extension length was calculated as the denoised and normalized distance between landmarks. (**d**) Using these data, we performed semi-automated detection of PE epochs by first identifying peaks from normalized proboscis extension lengths. Then we detected the start (cyan triangle) and end (magenta triangle) of these events. We removed false-positive detections by thresholding the amplitude (cyan line) and duration (magenta line) of events. Finally, we generated a binary trace of PE epochs (shaded area).

**Figure S3:**
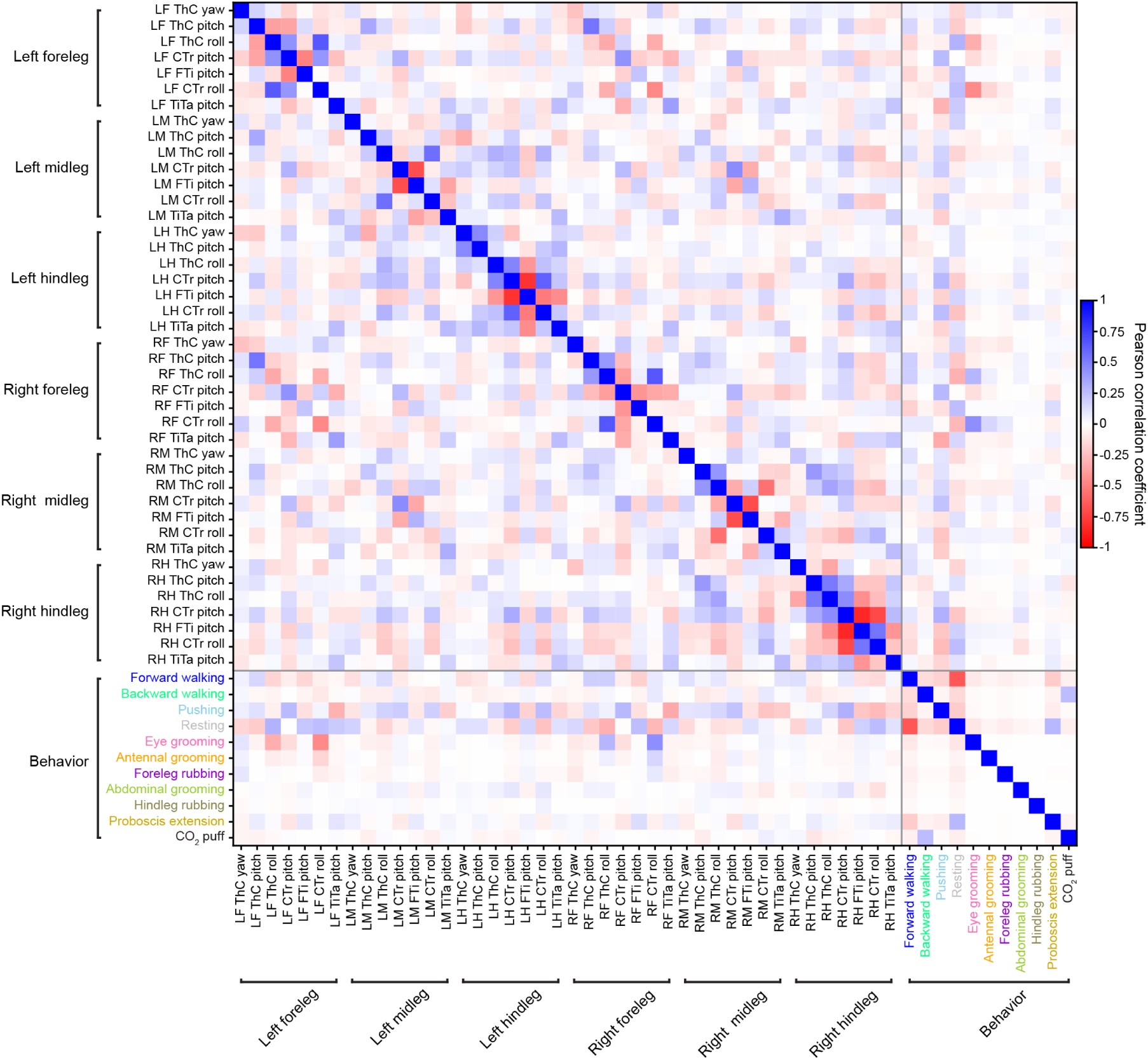
Correlations among and between low-level joint angles and high-level behaviors. Pearson correlation coefficients (color-coded) for joint angles, high-level behavioral states, proboscis extensions, and puffs.

**Figure S4:**
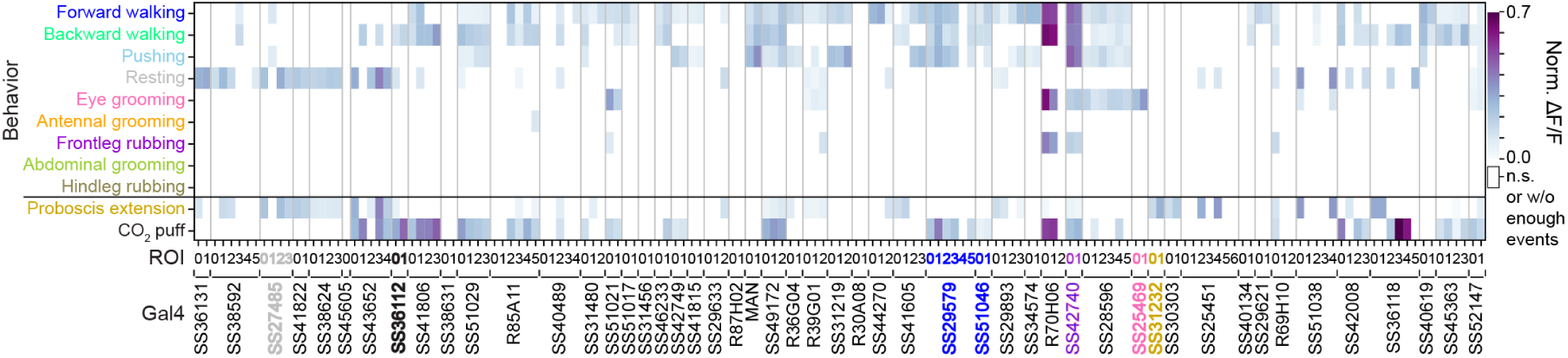
Normalized mean activity (Δ*F/F*) of ascending neurons during high-level behaviors. Normalized mean Δ*F/F* for a given AN across all epochs of a specific high-level behavior. Analyses were performed for 157 ANs recorded from 50 driver lines. Lines selected for more in-depth analysis are color-coded by the behavioral state best explaining their neural activity: SS27485 (resting), SS36112 (puff responses), SS29579 (walking), SS51046 (turning), SS42740 (foreleg-dependent behaviors), SS25469 (eye grooming), and SS31232 (proboscis extensions). Note that fluorescence for non-orthogonal behaviors/events may overlap (e.g., for backward walking and puff, or resting and proboscis extensions). To minimize contamination due to signals from preceding behaviors (resulting from the long decay kinetics of calcium indicators), conditions with less than ten epochs longer than 0.7 s are masked (white). Δ*F/F* signals are normalized between 0 and 1 to minimize the influence of differences in calcium indicator expression levels on data interpretation. ANOVA and posthoc Tukey tests to correct for multiple comparisons were performed to test if values are significantly different from baseline. Non-significant samples are also masked (white).

**Figure S5:**
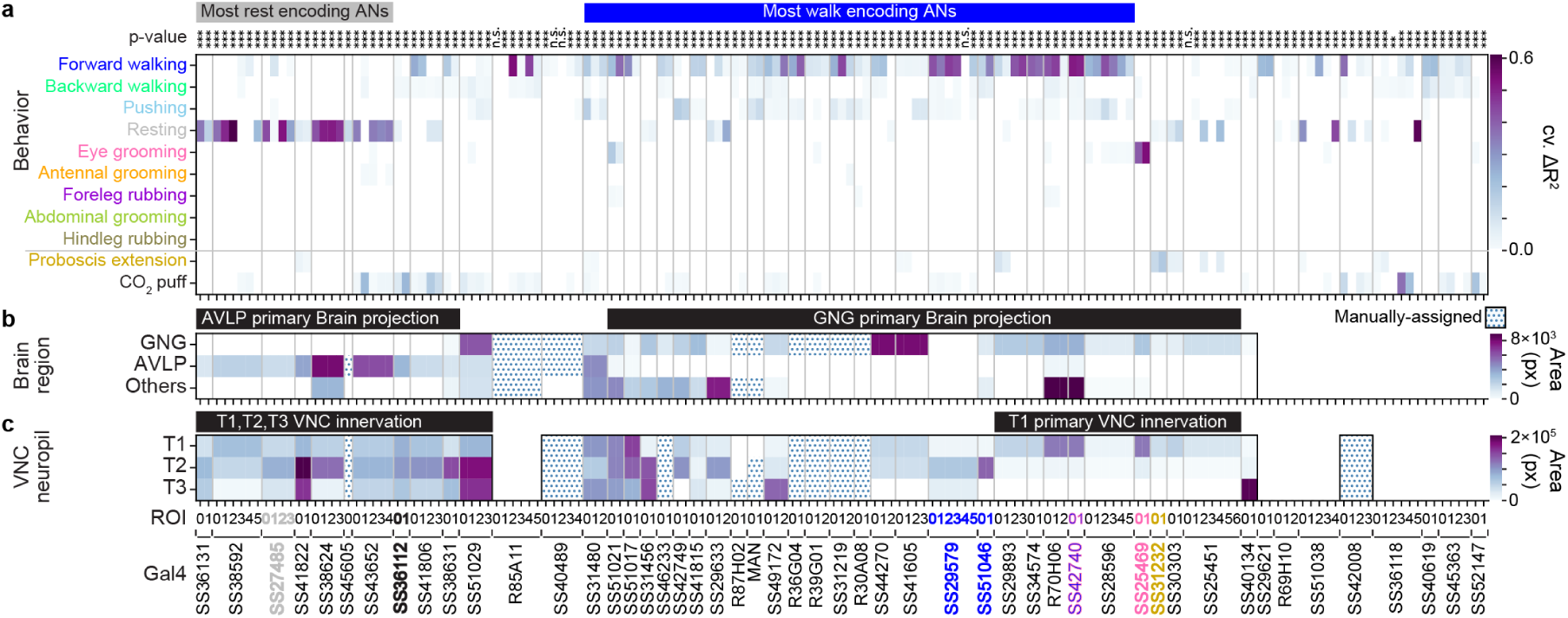
Relationship between ascending neuron behavioral encoding, brain targeting, and VNC patterning. **(a)** Variance in AN activity that can be uniquely explained by a regressor (cross-validated Δ*R*^2^) for high-level behaviors. Regression analyses were performed for 157 ANs recorded from 50 driver lines. Lines (and their corresponding ANs) selected for more in-depth analysis are color-coded by the behavioral class that best explains their neural activity: SS27485 (resting), SS36112 (puff responses), SS29579 (walking), SS51046 (turning), SS42740 (foreleg-dependent behaviors), SS25469 (eye grooming), and SS31232 (proboscis extensions). Non-orthogonal regressors (PE and CO_2_ puffs) are separated from the others. *P* -values report the F-statistic of overall significance of the complete regression model with no regressors shuffled (\**p<*0.05, \*\**p<*0.01, and \*\*\**p<*0.001). The most substantial AN **(b)** targeting of brain regions, or **(c)** patterning of VNC regions, as quantified by pixel-based analysis of MCFO labelling. Driver lines that were manually quantified are indicated (dotted cells). Projections that could not be unambiguously identified are left blank. Notable encoding and innervation patterns are indicated by bars above each matrix.

**Figure S6:**
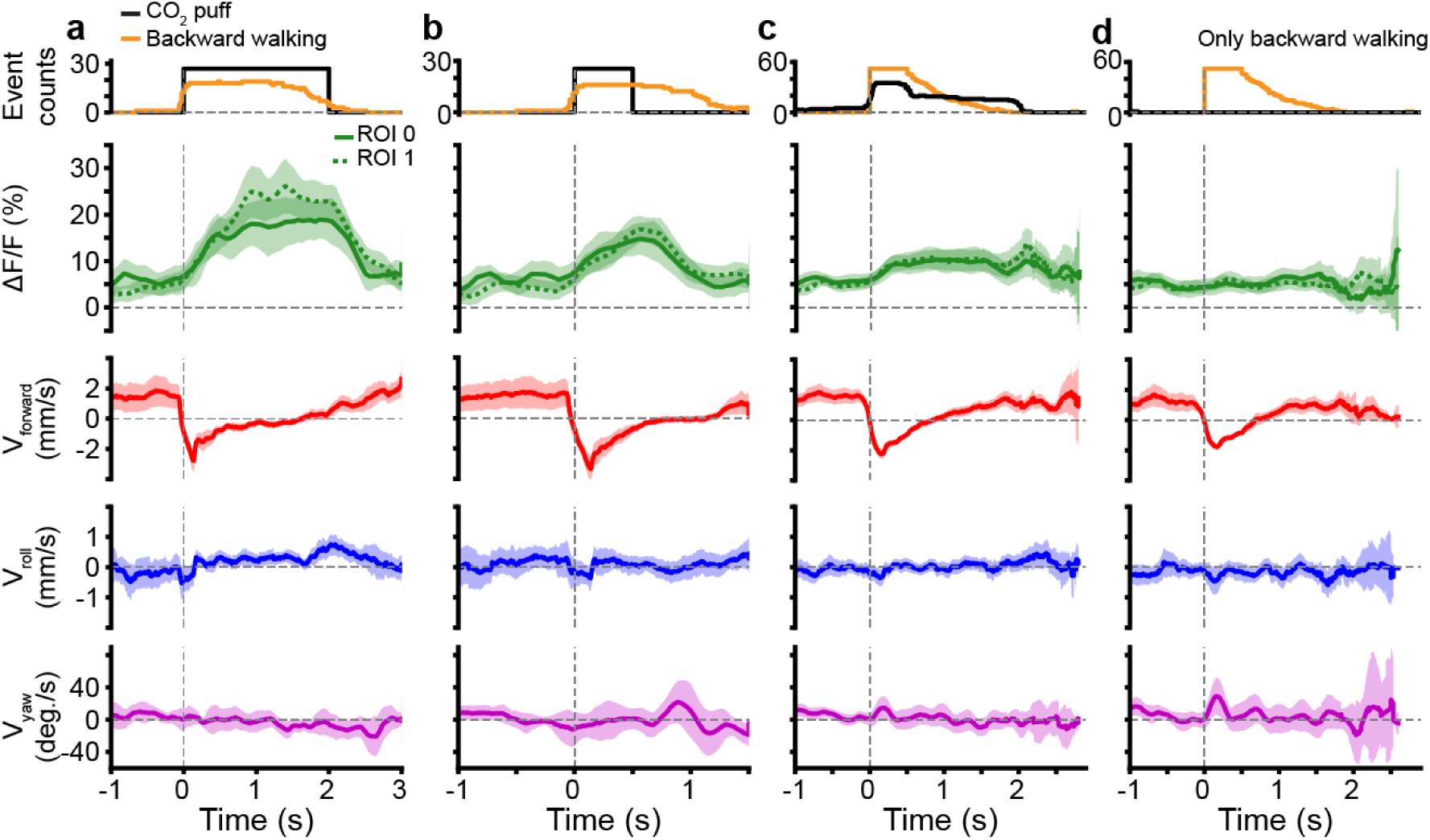
Puff-ANs respond to CO_2_ puffs and do not encode backward walking. Puff-ANs (SS36112) activity (green) and corresponding spherical treadmill rotational velocities (red, blue, and purple) during **(a)** long, 2 s CO_2_-puff stimulation (black) and associated backward walking (orange), **(b)** short, 0.5 s CO_2_-puff stimulation, **(c)** periods with backward walking, and **(d)** the same backward walking events as in **c** but only during periods without coincident puff stimulation. Shown are the mean (solid and dashed lines) and 95% confidence interval (shaded areas) of multiple Δ*F/F* and ball rotation time-series.

**Figure S7:**
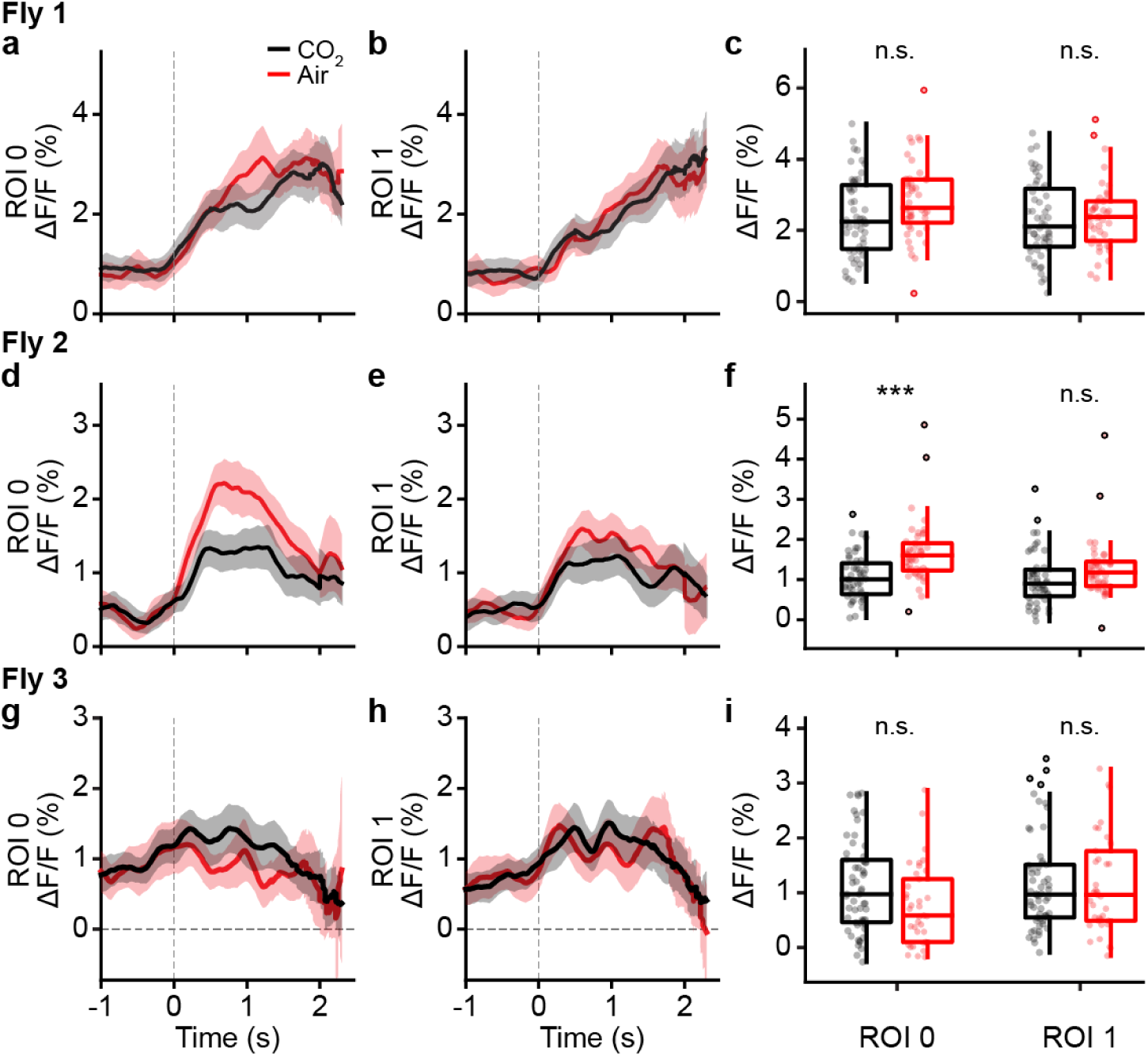
Puff-ANs respond similarly to puffs of air, or CO_2_. Activity of puff-ANs (SS36112) from three flies (**a-c**, **d-f**, and **g-i**, respectively) in response to puffs of air (red), or CO_2_ (black). **(a-b, d-e, g-h)** Shown are mean (solid and dashed lines) and 95% confidence interval (shaded areas) Δ*F/F* for ROIs **(a, d, g)** 0 and **(b, e, h)** 1. **(c, f, i)** Mean fluorescence (circles) of traces for ROIs 0 (left) or 1 (right) from 0.7 s after puff onset until the end of stimulation. Overlaid are box plots representing the median, interquartile range (IQR), and 1.5 IQR. Outliers beyond 1.5 IQR are indicated (opaque circles). A Mann-Whitney test (*** p*<*0.001, ** p*<*0.01, * p*<*0.05) was used to compare responses to puffs of CO_2_ (red), or air (black).

**Figure S8:**
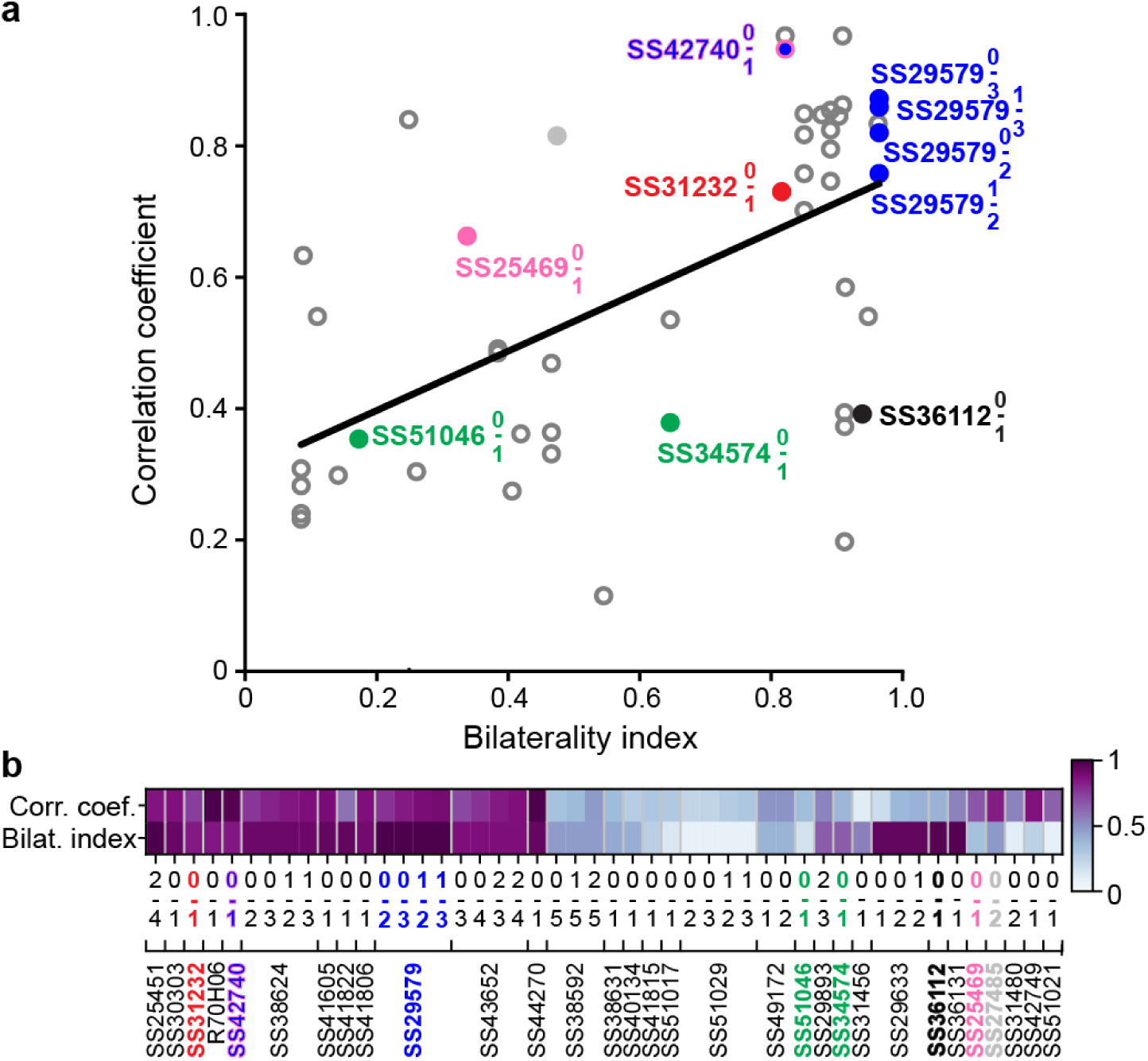
The bilaterality of an ascending neuron pair’s VNC patterning correlates with the degree of synchronous activity. **(a)** A bilaterality index, quantifying the differential innervation of left and right VNC (without distinguishing between axons and dendrites) is compared with the Pearson correlation coefficient for activity of left and right ANs within a pair (R^2^ = 0.31 and p*<*0.001 using an F-test). **(b)** Bilaterality index and Pearson correlation coefficient values for each AN pair examined.

**Figure S9:**
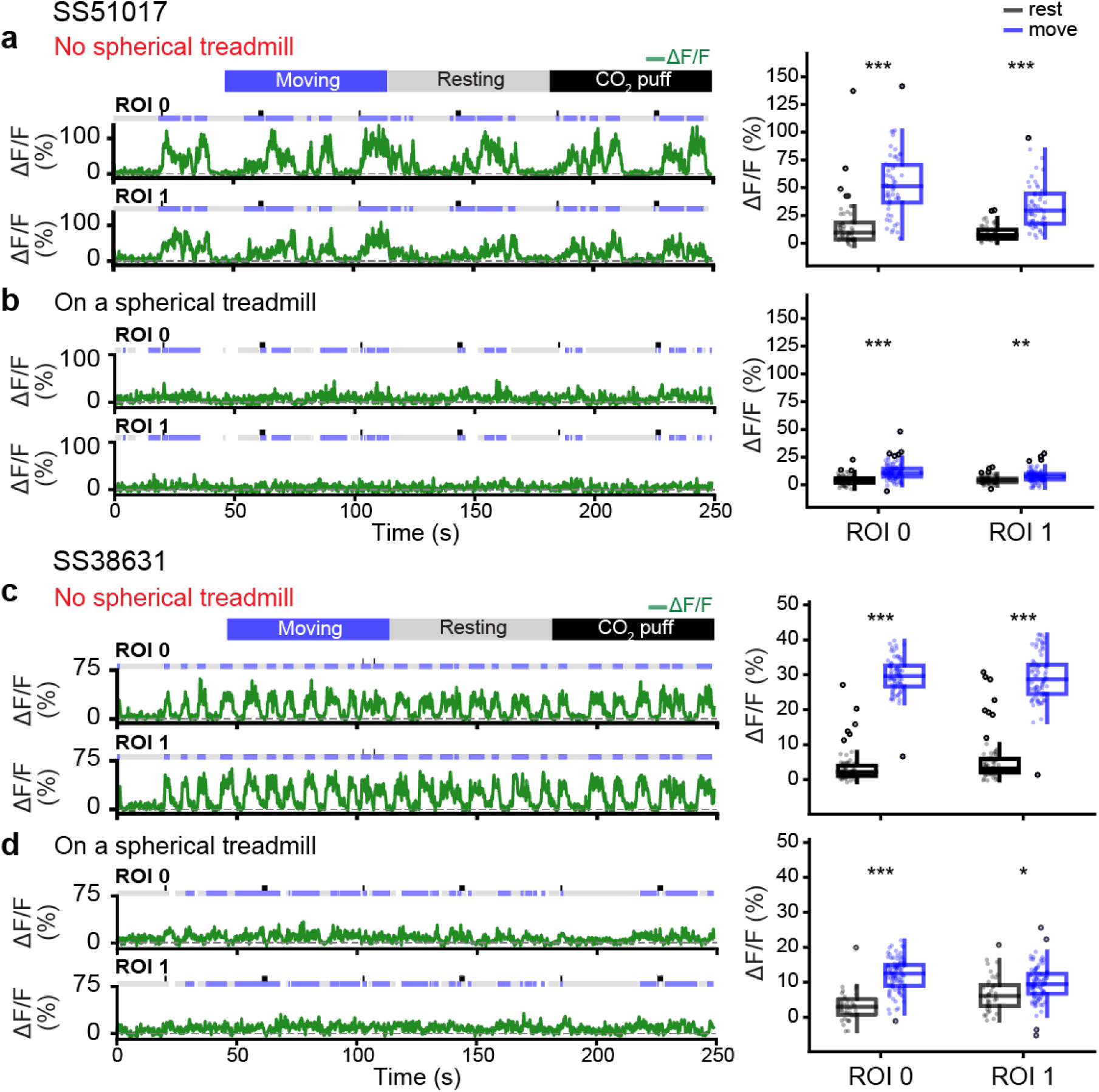
Ascending neurons that become active only in the absence of the spherical treadmill. Representative AN recordings from ROIs 0 and 1 for (**a, b**) one SS51017-spGal4 animal, or (**c, d**) one SS38631-spGal4 animal measured when it is (**a, c**) suspended without a spherical treadmill, or (**b, d**) in contact with the spherical treadmill. Moving, resting, and puff stimulation epochs are indicated. Shown are (**left**) representative neural activity traces and (**right**) summary data including the median, interquartile range (IQR), and 1.5 IQR of AN Δ*F/F* values when the animal are resting (black), or moving (blue). Outliers (values beyond 1.5 IQR) are indicated (black circles). Statistical comparisons were performed using a Mann-Whitney test (*** p*<*0.001, ** p*<*0.01, * p*<*0.05).

**Figure S10:**
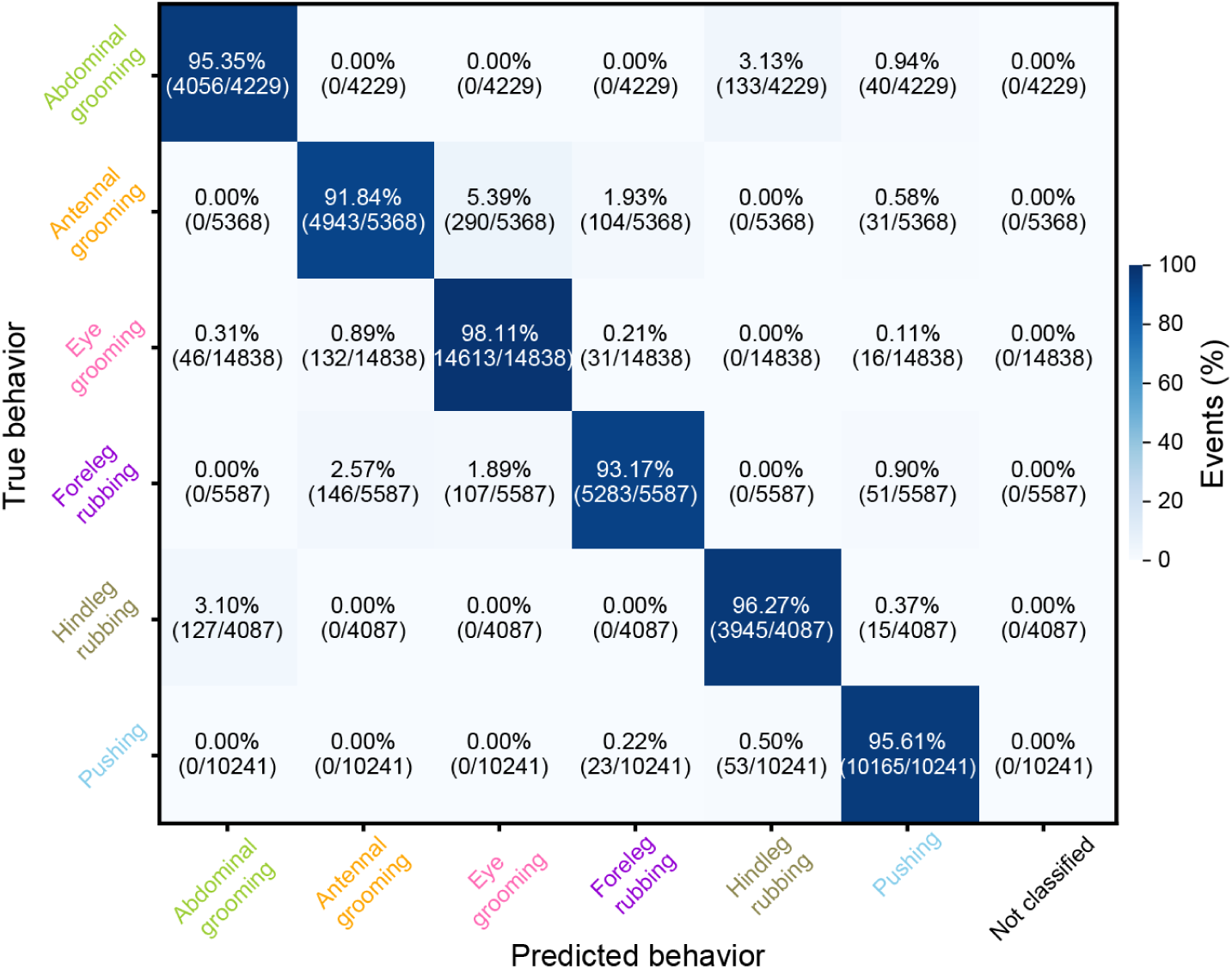
Behavior classifier accuracy. A confusion matrix quantifies the accuracy of predictions using 10-fold, stratified cross-validation of a histogram gradient boosting classifier. Walking and resting are not included in this evaluation because they are predicted using spherical treadmill rotation data. The percentage of events in each category (‘predicted’ behavior versus ground-truth, manually-labelled ‘true’ behavior) is color-coded.

**Figure S11:**
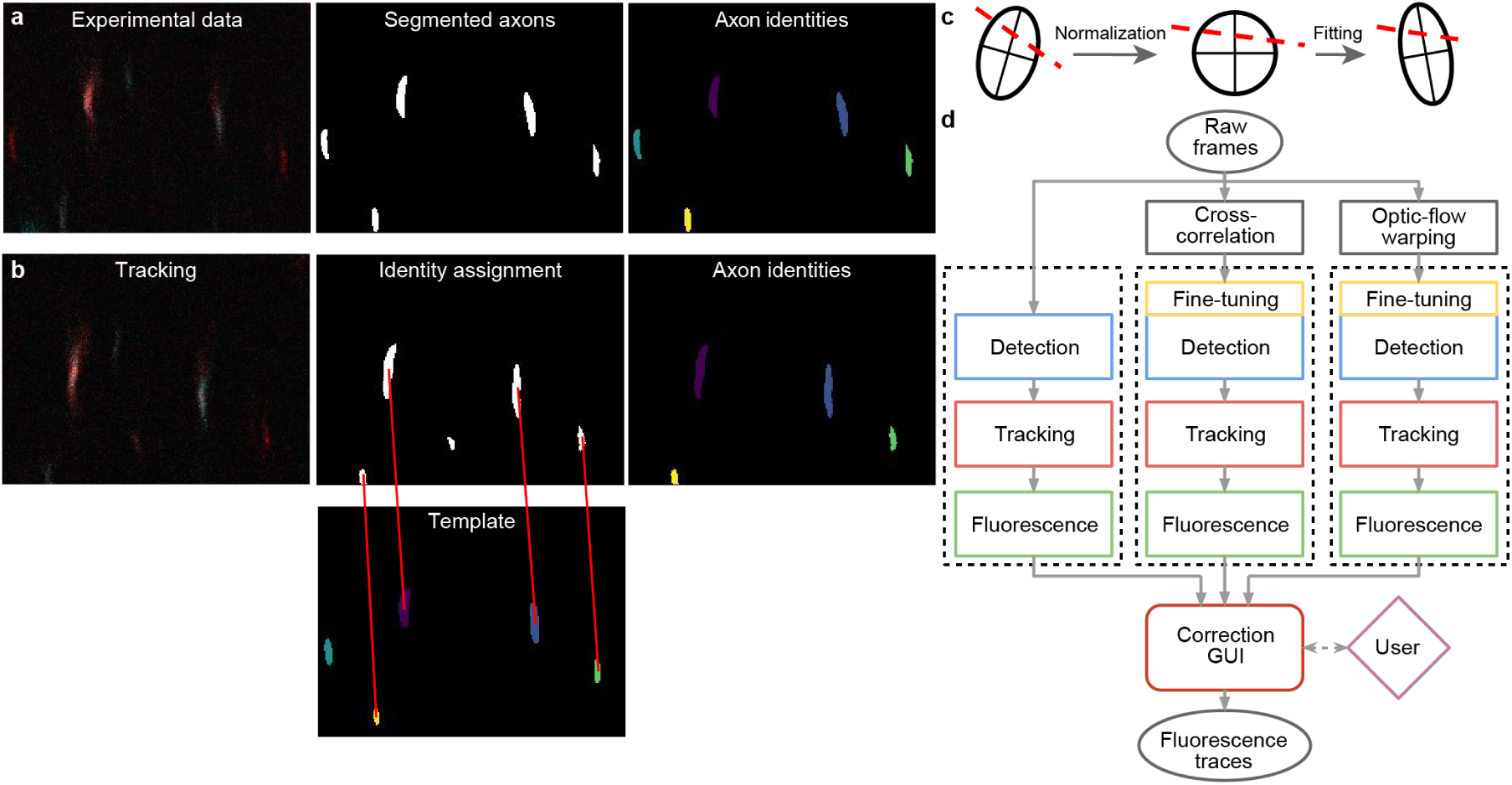
AxoID, a deep learning-based algorithm tracks axonal cross-sections in two-photon microscopy images. **(a)** Pipeline overview: a single image frame (left) is segmented (middle) during the detection stage with potential axons shown (white). Tracking identities (right) are then assigned to these ROIs. **(b)** To track ROIs across time, ROIs in a tracker template (bottom-middle) are matched (red lines) to ROIs in the current segmented frame (top-middle). An undetected axon in the tracker template (cyan) is left unmatched. **(c)** ROI separation is performed for fused axons. An ellipse is first fit to the ROI’s contour and a line is fit to the separation (dashed red line). For normalization, the ellipse is transformed into an axis-aligned circle and the linear separation is transformed accordingly. For another frame, a transformation of the circle into a newly fit ellipse is computed and applied to the line. The ellipse’s main axes are shown for clarity. **(d)** The AxoID workflow. Raw experimental data is first registered via cross-correlation and optic flow warping. Then, raw and registered data are separately processed by the fluorescence extraction pipeline (dashed rectangles). Finally, a GUI is used to select and correct the results.

## Notes

https://dataverse.harvard.edu/dataverse/AN

https://github.com/NeLy-EPFL/Ascending_neuron_screen_analysis_pipeline

https://github.com/NeLy-EPFL/AxoID

### Competing Interest Statement

The authors have declared no competing interest.

